# ARID1A-induced transcriptional reprogramming rewires signalling responses to drug treatment in melanoma

**DOI:** 10.1101/2024.12.05.626952

**Authors:** Charlie George Barker, Sumana Sharma, Ana Mafalda Santos, Konstantinos-Stylianos Nikolakopoulos, Athanassios D. Velentzas, Franziska I. Völlmy, Angeliki Minia, Vicky Pliaka, Maarten Altelaar, Gavin J Wright, Leonidas G. Alexopoulos, Dimitrios J. Stravopodis, Evangelia Petsalaki

**Affiliations:** European Molecular Biology Laboratory, European Bioinformatics Institute (EMBL-EBI), Wellcome Genome Campus, Hinxton, Cambridgeshire, CB10 1SD, United Kingdom; University College London Cancer Institute, London WC1E 6DD, UK; Radcliffe Department of Clinical Medicine and Medical Research Council, Translational Immune Discovery Unit, Weatherall Institute of Molecular Medicine, University of Oxford, Oxford, United Kingdom; Section of Cell Biology and Biophysics, Department of Biology, School of Science, National and Kapodistrian University of Athens (NKUA), 15701 Athens, Greece; Biomolecular Mass Spectrometry and Proteomics, Center for Biomolecular Research and Utrecht Institute for Pharmaceutical Sciences, Utrecht University, Utrecht, The Netherlands; Protavio, Demokritos Technology Park, Building 27, Patriarchou Grigoriou & Neapoleos 27, 15341 Aghia Paraskevi, Attiki, Greece; Netherlands Proteomics Center, Padualaan 8, 3584 CH Utrecht, The Netherlands Mass Spectrometry and Proteomics Facility, The Netherlands Cancer Institute, 1066 CX Amsterdam, The Netherlands; Department of Biology, Hull York Medical School, York Biomedical Research Institute, University of York, York, UK; Department of Mechanical Engineering, National Technical University of Athens, Athens, Greece

## Abstract

Resistance to BRAF and MAPK inhibitors is a significant challenge in melanoma treatment, driven by adaptive and acquired mechanisms that allow tumour cells to evade therapy. Here, we examined early signalling responses to single and combined BRAF and MAPK inhibition in a BRAFV600E, drug-sensitive melanoma cell line and a drug-resistant ARID1A-knockout (KO) derivative. ARID1A, frequently mutated in melanoma, is associated with resistance and immune evasion. Using an innovative systems biology approach that integrates transcriptomics, proteomics, phosphoproteomics, and functional kinomics through matrix factorization and network analysis, we identified key signalling alterations and resistance mechanisms.

We found that ARID1A-KO cells exhibited transcriptional rewiring, sustaining MAPK1/3 and JNK activity post-treatment, bypassing feedback sensitivity observed in parental cells. This rewiring suppressed PRKD1 activation, increased JUN activity—a central resistance network node—and disrupted PKC dynamics through elevated basal RTKs (e.g., EGFR, ROS1) and Ephrin receptor activity post-treatment. ARID1A mutations also reduced HLA-related protein expression and enriched extracellular matrix components, potentially limiting immune infiltration and reducing immunotherapy efficacy. Our graph-theoretical multi-omics approach uncovered novel resistance-associated signalling pathways, identifying PRKD1, JUN, and NCK1 as critical nodes. While receptor activation redundancies complicate single-target therapies, they also present opportunities for combination strategies.

This study highlights ARID1A’s role in reshaping signalling and immune interactions, offering new insights into melanoma resistance mechanisms. By identifying actionable targets, including JUN and immune pathways, we provide a foundation for developing integrated therapeutic strategies to overcome resistance in BRAF/MAPK inhibitor-treated melanoma.

**One sentence summary:** This study reveals how ARID1A-mediated transcriptional rewiring drives resistance to MAPK inhibitors in melanoma by altering signalling pathways, immune interactions, and receptor dynamics, highlighting potential targets for combinatorial therapies.

## Introduction

Melanoma is an aggressive form of skin cancer arising from melanocytes. It is largely driven by aberrant cellular signalling processes, specifically in the mitogen-activated protein kinase (MAPK) pathway, with nearly 40-50% of all melanomas harbouring mutations in the central MAPK pathway kinase BRAF (*1*). The second most common mutation accounting for approximately 30% of all melanomas is in neuroblastoma RAS viral oncogene homolog (NRAS), an upstream kinase regulator of the MAPK pathway. Mutated BRAF, which in 80% of all BRAF mutations is BRAFV600E, is constitutively active and phosphorylates MEK proteins (MEK1 and MEK2), which in turn activate the downstream MAP kinases and aberrant cell proliferation. Melanoma cells harbouring mutated BRAF typically exhibit a dependency on the BRAF protein and the components of the MAPK pathway (*2*). BRAF inhibitors alone produce highly effective outcomes initially; however, these effects are short-lived, as resistance mechanisms frequently emerge, leading to the reactivation of the MAPK pathway and the development of adverse cutaneous effects (*3*). Combination therapies were developed to counter this by also inhibiting MEK, which led to longer progression-free survival (PFS) (*4*). Pharmacological inhibitors designed to target the mutated BRAF, such as vemurafenib and dabrafenib, in combination with MEK inhibitors, particularly trametinib, have become standard treatments in clinical settings for melanoma, specifically for patients with activating BRAF mutations (*5*).

Despite their effectiveness, the response duration to these treatments are still short-lived, and resistance develops in the majority of patients (*6*). Studies suggest that around 50% of patients treated with BRAF or MEK inhibitors experience disease relapse and progression within 6 to 7 months of initiating treatment (*7*). Resistance to BRAF inhibitors in around 80% of the cases involves genetic and epigenetic changes, leading to the reactivation of the MAPK pathway through the ‘re-wiring’ of cellular signalling processes (*6*, *8–14*). Other mechanisms include reactivation of PI3K-mTOR pathway through inactivation of PTEN phosphatase or via stromal cells within the tumour microenvironment that secrete growth factors to activate receptor tyrosine kinases (RTKs) of both PI3K and MAPK pathways (*15–17*). Additionally, it has also been shown that cells can develop resistance without acquiring new mutations but rather by temporary and reversible adaptations to selective pressure (*18*, *19*). Studies have described the existence of persister cells, i.e., cells that continue to grow in culture for a long time even if the oncogenic BRAF is inhibited in culture (*20*). How adaptive changes in cells relate to acquired resistance is still not fully understood, with some studies suggesting that cells that have undergone adaptive resistance might have low accuracy of DNA replication and low efficacy of DNA damage responses compared to drug-naïve cells leading to the accumulation of resistance mutations (*18*, *21*). A clear understanding of both how intracellular signalling is immediately rewired upon perturbation and how more stable resistance is achieved over the long term is necessary to design future therapies that are more efficacious and longer lasting.

As an alternative to targeted therapy, immunotherapy approaches alleviate the inhibition of the immune system by blocking inhibitory receptors allowing immune cells to eliminate cancer cells (*22*). In melanoma, inhibitory checkpoint blockers, particularly anti-programmed cell-death protein 1 (PD-1) and anti-cytotoxic T-lymphocyte-associated antigen 4 (CTLA-4), have demonstrated durable responses in a subset of patients. While these responses tend to be more enduring, the overall response rate to immunotherapy is relatively modest, estimated at 40-50% of patients (*23*). Consequently, current recommendations advocate for the use of both targeted therapies and immunotherapies as the first-line treatment for metastatic melanoma (*23*),(*24*).

Targeted therapy in melanoma is based on the dependency of the cells on the mutated pathway. These include hotspot mutations in genes such as *BRAF*, *NRAS*, *KIT*, *GNAQ* and others. However, there is one frequently mutated gene in melanoma; the one encoding the AT-Rich Interaction Domain 1A (ARID1A) protein, which stands out as it is mutated without a distinct hotspot, which tends to be a pattern for tumour suppressor genes (*25*). ARID1A is a key component of the switching/sucrose nonfermentable (SWI/SNF) complex. This complex is known for its pivotal role in chromatin remodelling and influencing tumour epigenetics. Approximately 11.5% of melanoma patients exhibit mutations in the *ARID1A* gene (*26*). Mutations in ARID1A are associated with elevated programmed cell death-ligand 1 (PD-L1) expression, a heightened tumour mutational burden (TMB), decreased infiltration of immune cells into the tumour microenvironment (TME), and compromised mismatch repair (MMR) (*27–30*). Notably, a number of genome-wide loss-of-function CRISPR screens have also identified *ARID1A* as a critical factor in conferring resistance to BRAF/MEK inhibitors like vemurafenib, and selumetinib (*31*, *32*). Despite these significant findings, the clinical significance of ARID1A mutations, particularly in the context of melanoma, remains ambiguous. As ARID1A has been implicated not only in resistance to targeted therapies for melanoma but also for modulating the therapeutic responses to immune checkpoint blockade, its study lends to better understanding of interplay between these mechanisms in developing resistance.

Here, we present an integrative multi-omics study to compare the response of drug sensitive ARID1A WT (wild-type) vs resistant ARID1A KO (knockout) melanoma cell lines to both single and combination drug perturbation. We devised a computational strategy that combined data integration using multi-omics factor analysis (MOFA (*33*)), with a network propagation-based method, phuEGO (*34*), to extract the predominantly affected signalling networks post-short-term treatment of both resistant and sensitive melanoma cell lines with BRAF inhibitors alone or in combination. This allowed us to interpret the data in a unified framework providing new insights into the signalling processes activated in response to single/combination drug treatment of both sensitive and resistant melanoma cells.

## Results

### Multi-omics data integration identifies molecular signatures associated with drug response and ARID1A KO

ARID1A has been previously identified as a hit in genome-wide screens that have been carried out to identify candidates that confer resistance to MAPK inhibitor (selumetinib) or BRAF inhibitor (vemurafenib) (*31*, *32*). We performed two additional genome-wide screens using A375 cells to identify genes required for resistance of these cells to another MAPK inhibitor (trametinib) and identified ARID1A in both replicates together with other genes that are often required for drug resistance in BRAFV600E mutant melanoma cells (e.g., *NF1/2*, *KIRREL*, *MED12*, *TAF5/6L*) suggesting that ARID1A plays a role in resistance to melanoma cells with BRAFV600E mutation in response to MAPK or BRAF inhibitor drugs (**Fig. 1A, Table S1**). We then used sgRNAs to target *ARID1A* and *MED12*, a previously characterised gene responsible for drug resistance to vemurafenib and treated the cells with trametinib for 6 hours. After this time point, we measured the level of phosphorylation of pMEK and pERK using a Luminex assay (*35*) and noted an increase in both pERK and pMEK phosphorylation in both mutants compared to ‘empty’ sgRNA transduced cell lines. Unlike in the ‘empty’ transduced cell line, the level of pERK in both mutants remained unchanged when the cells were treated with trametinib suggesting that these cells were non-responsive to MAPK inhibition (**Fig. 1B**).

**Figure 1.**
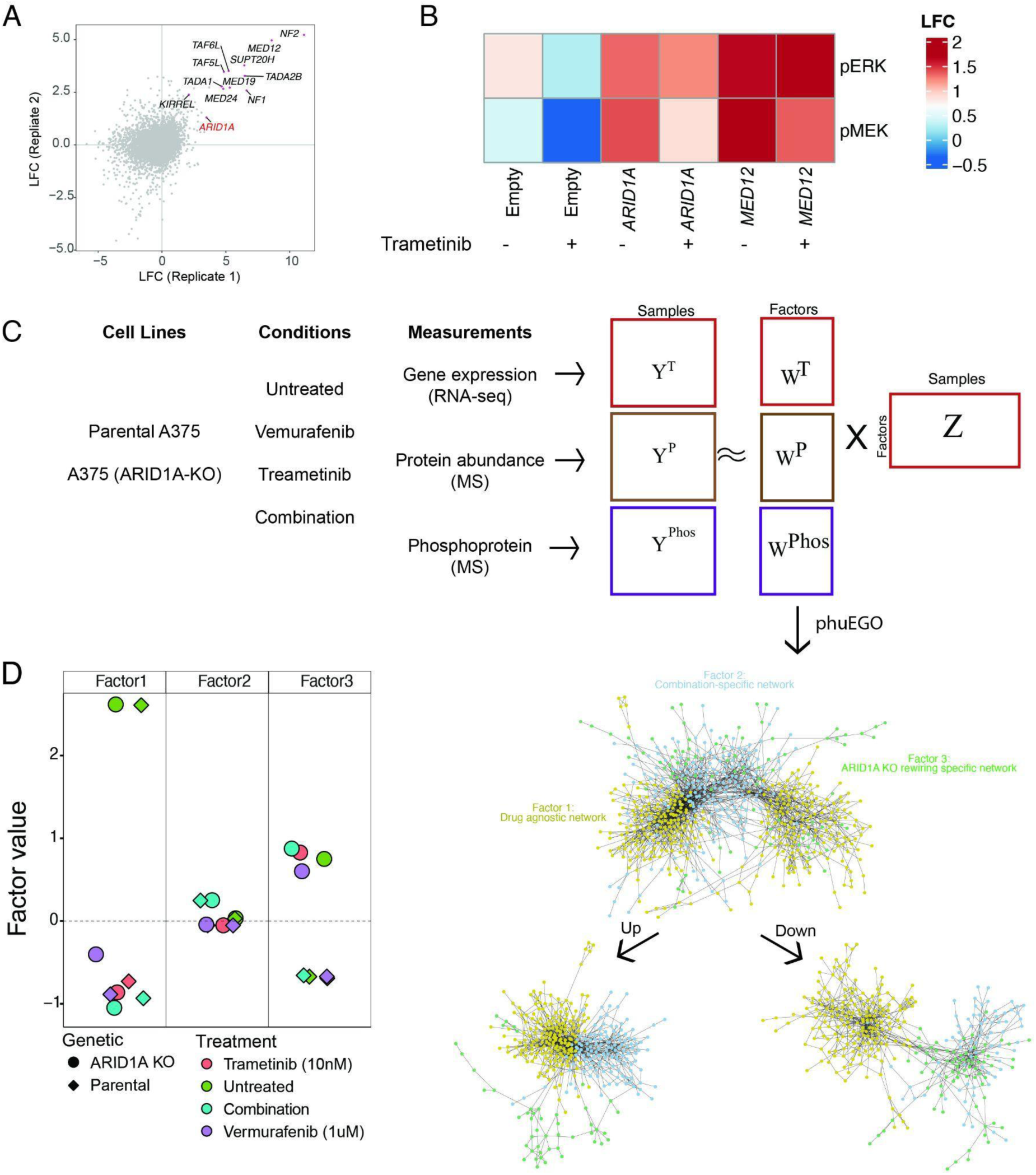
Overview of study design. **A.** ARID1A KO confers a survival advantage in a pooled genome-wide CRISPR/Cas9 screens of A375 cells treated with trametinib (MEKi). **B.** ARID1A KO cells do not show reduced phosphorylation of ERK upon treatment with trametinib **C.** Schematic presentation of the method employed to integrate and reconstruct signalling networks from melanoma multi-omics data. Up and down indicate positive and negative weights in the factors. **D.** Factor weight loadings (y axis) for the different samples (colours for drug treatment and shapes for genetic conditions).

To study the differences between signalling responses and gene regulation to single and combination drug treatment in sensitive versus resistant ARID1A-knockout (KO) cell lines, we acquired an early passage parental A375 cell line and a matched ARID1A KO line (**Fig. S1A and B**). This cell line exhibited increased resistance to both vemurafenib and trametinib treatment (**Fig. S1C**). To comprehensively characterise the signalling network rewiring underpinning the resistance of ARID1A KO to BRAF/MEK inhibition, we collected multi-omics data from parental and ARID1A KO cells in the presence or absence of drugs. Specifically, we collected mass spectrometry-based proteomics, phosphoproteomics as well as transcriptomics data upon no treatment or treatment with either trametinib, or vemurafenib, or both drugs for 6 hours (**Materials and Methods; Fig. 1C**). We used 6 hours as a time point, as this provided a single steady-state measurement post-treatment of drugs and our pilot measurement of phosphosites on 17-plex luminex assay showed that the effect of MAPK inhibitor on suppression of phosphorylation of key signalling proteins was intact at this time point (**Fig. S1D**).

From the mass spectrometry we quantified 8,139 proteins and 3,207 phosphosites after integration of our different experimental runs (**Materials and Methods; Table S2 and S3**). Using RNAseq we quantified the transcription of 14,376 genes (**Table S4**). All datasets were reproducible (**Fig. S2A-C**). Among the 715 proteins, 372 phosphopeptides and 7,557 genes that were (significantly) differentially abundant (FDR adjusted p value < 0.01) between WT vs ARID1A KO experiments (**Fig. S3; Tables S2-4**), only 13 were common to all datasets demonstrating the orthogonal nature of the different omics layers (**Fig. S3B**).

We next used multi-omics factor analysis (MOFA (*33*)) to identify latent factors that explain the variation of all modes of data in a prior knowledge-agnostic manner (**Fig. 1C**). Our analysis revealed 3 factors describing the adaptive response to drug treatments (Factors 1 and 2) and the sustained resistance response illustrating the effect of the loss of ARID1A (Factor 3) (**Fig. 1D**). The low-dimensional representation of the data illustrated that the differences between the isolated drug treatments (vemurafenib and trametinib) were marginal at the level of the omics data (**Fig. S2A; Table S5**) compared to the drug combination treatment.

Variance decomposition showed that the drug associated factors explain the majority of variance in both protein and phosphosite, and mRNA abundance (**Fig. S4**). Factor 2 (describing the differences that characterise combination therapy), explains no variance in the transcriptome (0%), but the majority of the variation is found in the phosphoproteomic data (32.5%) and to a lesser extent the proteomic data (28.5%). Factor 3 (which is associated with the ARID1A KO) appears to describe variance in the transcriptome (12.7%) and proteome (4.3%) and to a much lesser extent the phosphoproteome (0.5%; **Fig. S4**).

To understand the functional implications on the signalling processes represented by the factors identified above, we sought to place the identified genes within the context of their functional environment, i.e., their interaction networks. To this end, we adapted phuEGO (*34*), a network propagation-based method, to extract active signatures from phosphoproteomic datasets (**Materials and Methods**), for use with the factor loadings taken from MOFA. PhuEGO combines network propagation with ego network decomposition allowing the identification of small networks that comprise the most functionally and topologically similar nodes to the input ones. This allowed us to generate minimal networks from the factor-specific loadings covering the proteins driving the differences between drug responses and the ARID1A KO, as a function of phosphoproteomic, transcriptomic and proteomic weights (**Fig. 1C)**. This was performed on upregulated and downregulated and then merged to produce 3 networks (**Data S1**).

### Drug-agnostic changes associated with treatment involve negative feedback of RTKs and MAPK

As mentioned above, the changes observed in factor 1 refer to those agnostics to the specific treatment, i.e., regardless of whether BRAF, MEK1/2 or both were inhibited. The most central nodes of the network included several receptors, kinases and transcription factors, with EGF, FYN, IGF1R, TEK, MAPK1 and MAPK3 followed by PRKD1, JUN, PTK2 and INSR being characteristic examples (**Fig. 2A and Fig. S5A**). Pathway enrichment analysis of the factor 1-associated network finds pathways related to ‘Melanoma’ and various terms related to MAPK signalling, including the term ‘DUSP regulation of MAPK pathway’ (**Fig. 2B and C**).

**Figure 2.**
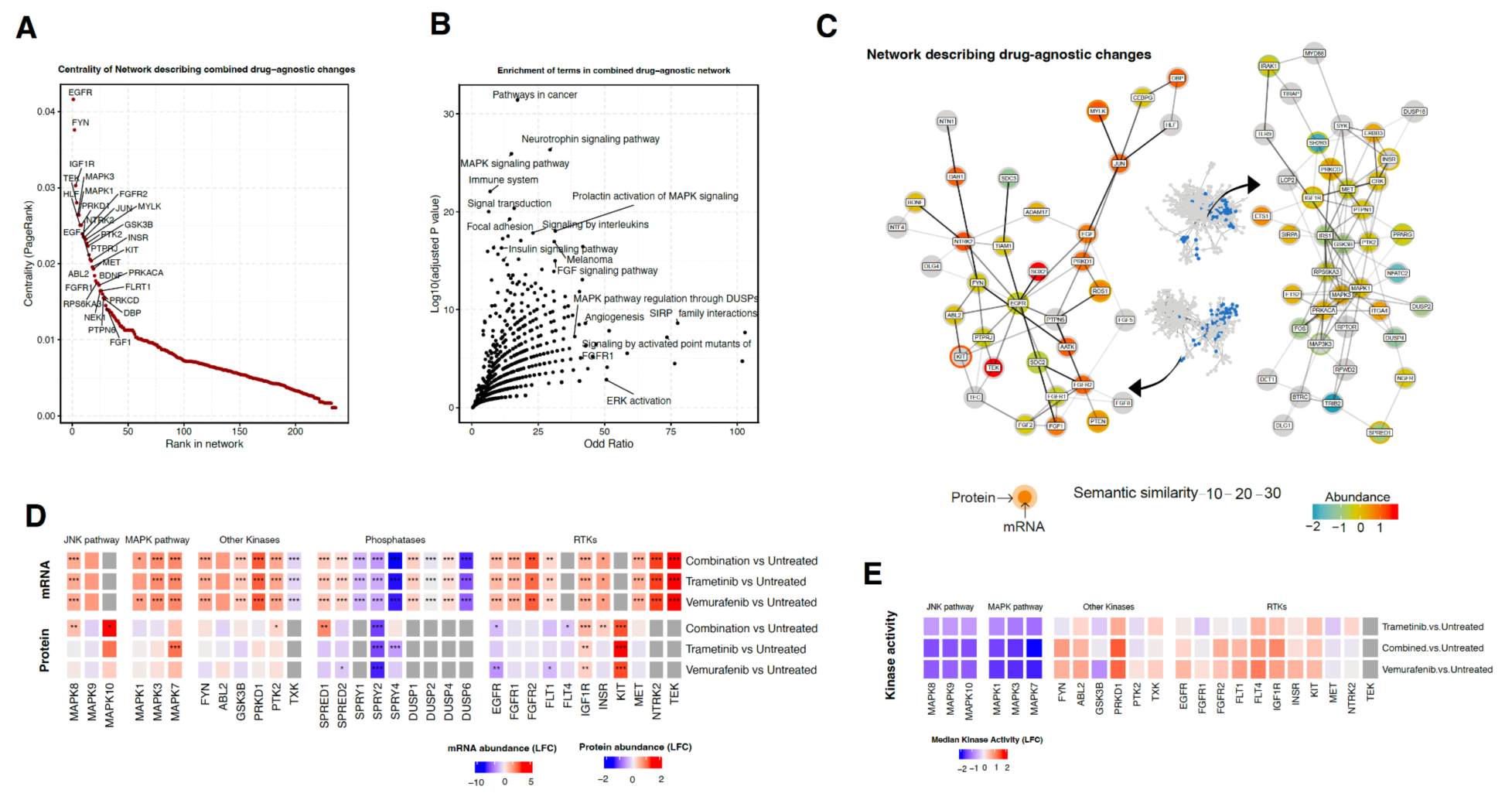
Overview of molecular signature associated with drug response regardless of type of drug (Factor 1). **A.** Nodes ordered (x axis) by their centrality (y axis) in the phuEGO-derived network that are associated with drug-agnostic responses. **B.** Processes significantly (y axis) enriched (x axis) in the drug-agnostic phuEGO-derived network. **C.** Subset of drug-agnostic phuEGO-derived network highlighting optimised to the 50 most central nodes and their interactions. **D.** Heatmap demonstrating the changes in proteomics and transcriptomics abundances for negative regulators of the RTK/MAPK pathways. **E.** Heatmap demonstrating the changes in proteomics and transcriptomics abundances, and functional kinomics measurements for multiple kinases found as regulated or relevant in the network derived from Factor 1.

In agreement with other studies (*36*), we observed the decrease in abundance of the known negative feedback regulators of MAPK (DUSP1/2/4), shown to interact with MAPK3 (ERK1) (**Fig. 2D and Fig. S5B**). In both SPRY1/2/4 and SPRED1/2, we detected a decrease in both RNA abundance and protein abundance (**Fig. 2D and Fig. S5B; Table S3 and S4**). The decrease of these negative regulators of RTK and MAPK signalling seems able to relieve downstream inhibition, leading to increased growth factor signalling and ameliorating the effect of MAPK inhibition by vemurafenib or trametinib.

As protein/transcriptome abundance does not correlate well with kinase activity and we only found a few phosphosites modulated, we also collected functional kinomics data using the PamChip technology (*37*), which provides an estimate of multiplex kinase activities in a cell lysate over an array of immobilized target-peptides (**Fig. S5C; Table S6**). The magnitude of changes in the activity of kinases, in this dataset, correlates strongly with the centrality of the kinases within our network, with MAPK1 and PRKD1 being the most central kinases quantified in our network (**Fig. S5D**). Among them, we found a strong reduction in the MAP kinase activities upon drug treatment (**Fig. 2E**), even though this was not mirrored by changes in their respective abundance. We also found several other activated kinases, including PRKD1, FYN and IGFR1, which were shown to be central in our network, as well as ABL2, FLT1 and FGFR2. EGFR presented only a very small increase in activity, which contrasts with its reduction in both transcriptomic and proteomics abundance. Taken together, these results indicate rewiring of RTK-driven signalling following drug perturbation.

### Combination therapy invokes phosphorylation patterns associated with DNA damage repair

Factor 2 illustrates changes that are associated with response to drug combination treatment (**Fig. 1D**) and are mostly derived from the phosphoproteomics data (**Fig. S4**). Drug combination treatments targeting both BRAF and MEK are currently the standard of care for patients with metastatic and non-resectable melanoma, and have shown longer remission of the disease in patients (*38*). Our functional kinomics data indicate that there is practically no difference in the effect on MAP kinases between the combination treatment and single treatment with vemurafenib (**Fig. 2E**). We also did not observe a difference in killing efficiency of combination treatment compared to mono-treatment for this cell line (**Fig. S1C**). We, therefore, decided to zoom into this factor to shed light on the phosphoproteomic differences observed for this level.

Applying phuEGO to the weights from Factor 2, maps the unique protein networks that are affected by combination therapy (**Fig. 1C; Fig. S6A**). We detect transmembrane receptors, specifically ERBB2, INSR and EGFR, being central in the network, and surrounded by differentially phosphorylated proteins (**Fig. 3A and B, and Fig. S6A**). TMEM30A, a protein involved in cell migration, is the second most central node followed by SRC. AKT1, an alternate growth regulator from MAPK, is also highly central (**Fig. 3A and B, and Fig. S6B**). The corresponding combination therapy-specific downregulated network is centred around the transcription factor MYC and the DNA damage response protein TP53BP1 (**Fig. S6B**). Also present are the RAF1 and KRAS signal transducers. MAP3K7 (TAK1) is also observed, as it has negative weights due to reduced phosphorylation (**Fig. 3B; Fig. S6C**) following combined inhibition of MAP2K1 and BRAF.

**Figure 3.**
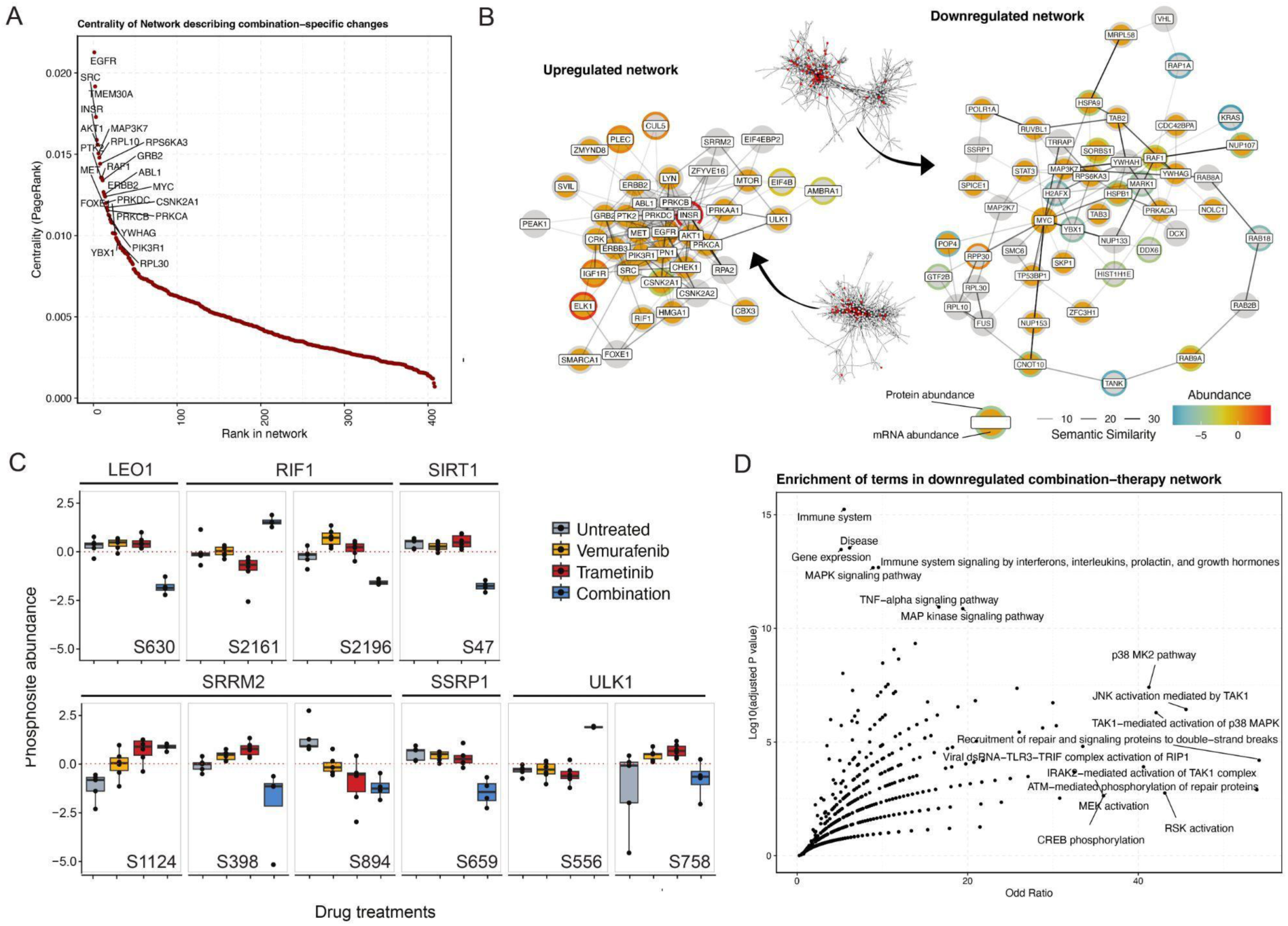
Overview of molecular signature associated specifically with combination drug response (Factor 2). **A.** Nodes ordered (x axis) by their centrality (y axis) in the phuEGO-derived network that are associated with drug-agnostic responses. **B.** Subset of combination treatment phuEGO-derived network highlighting the 30 most central nodes and their interactions. **C.** Highlighted phosphosite changes in the combination treatment showing the change in abundance (y axis) after treatment with drugs (x axis). **D.** Processes significantly (y axis) enriched (x axis) in the downregulated network upon combination treatment.

Despite phosphoproteomics being the main driver for this factor, there was little change in kinase activities, when looking at the functional kinomics data, compared to changes with the single-drug perturbations (**Fig. S6C**). Looking at the phosphosites that drive the variance captured by Factor 2, we find several phosphoproteins involved in DNA repair-related functions (**Fig. 3C** and **Table S5**). For example, RIF1 is known to be a key regulator of TP53BP1 able to promote non homologous end joining (NHEJ) DNA repair of double strand DNA breaks (*39*). We found several phosphosites being regulated in both proteins. While there are no functional annotations for these sites, RIF1 - S2196, which is downregulated in the combination treatment compared to no or single-drug treatment (**Fig. 3C**) is very close to S2205, which is known to inhibit protein’s function (*40*) and is predicted to be phosphorylated by JNK1,3 or P38δ or **γ**, among other kinases, all of which are downregulated in our functional kinomics dataset compared to no drug treatment (*41*). SSRP1, which is also known to be involved in DNA repair processes (*42*), shows decreased phosphorylation in S659 (**Fig. 3C**). ULK1-S556, which is known to be phosphorylated by ATM (*43*) and to be inducing autophagy (*44*), shows strong upregulation in the combination treatment compared to all other conditions (**Fig. 3C)**. SIRT1-S47 is not one of the phosphosites driving factor 2, however it is one of the significantly downregulated peptides in combination treatment versus untreated control, and is known to promote epithelial-to-mesenchymal transition (EMT) through autophagic degradation of E-cadherin (*45*). LEO1 - S630 is involved in the maintenance of embryonic stem cell pluripotency (*46*, *47*) and is also downregulated. Finally, we observe phosphoregulation of SRRM2, a component of spliceosome, at 3 phosphosites, with 2 of them being downregulated and one upregulated (**Fig. 3C)**.

Zooming out and to look at the processes involved in this network, we performed functional enrichment analysis on both up- and downregulated networks. In the upregulated network (**Fig. S6D**), we see significantly enriched terms corresponding to RTK-driven signalling (ERBB signalling pathway, ERBB2 signalling pathway and cMET signalling pathway). We also observe terms associated with PI3K/Akt signalling and mTOR signalling (‘PI3K events in ERBB2 signalling’ and ‘mTOR signalling pathway’). In the downregulated network, we see terms consistent with the expected inhibition of MAPK signalling, such as ‘MAPK signalling pathway’. We also pinpoint terms associated with DNA damage repair (‘ATM-mediated phosphorylation of repair proteins’ and ‘Recruitment of repair and signalling proteins to double-strand breaks’), as well as processes associated with the immune system and associated signalling (TNF-alpha, Interferons, Interleukins, Prolactin and Growth hormone; **Fig. 3D**).

### The basal transcriptional state of cells following ARID1A KO influences response to drug treatment

Even though the ARID1A-KO cell line is resistant to MEK/BRAF inhibition (**Fig. 4A**), the general networks associated with drug response appear to be very similar at the transcriptome and proteome levels (**Fig. 1D; Fig. S7A and S7B**). At the multiplex kinase activity level, however, we observe distinct differences, such as MAPK1/3’s insensitivity to drug treatment, switching activity of PRKD1 and FYN in response to treatments and overall increase in activity of the Ephrin receptor family **(Fig. 4A)**. If both conditions respond to drug treatment using the same molecular machinery (as represented by the ‘omics’), how can there be different responses and non-responsiveness to drug treatment at the signalling level? Looking at the initial state of the ARID1A KO cell lines, we observed differences in the initial abundances of several receptor tyrosine kinases (RTKs). These include receptors also increasing in the response to drug therapy, including ROS1 and ITGA4 (**Fig. 4B**). Other receptors are increasing in abundance, including EGFR (**Fig. 4C**) and CD44, while NGFR and IL6R are decreasing in abundance. Oncogenic transcription factors JUN and MYC are increasing in abundance, and so are their corresponding regulons (**Fig. S7C**), indicating an increase in activity and oncogenesis following ARID1A KO.

**Figure 4.**
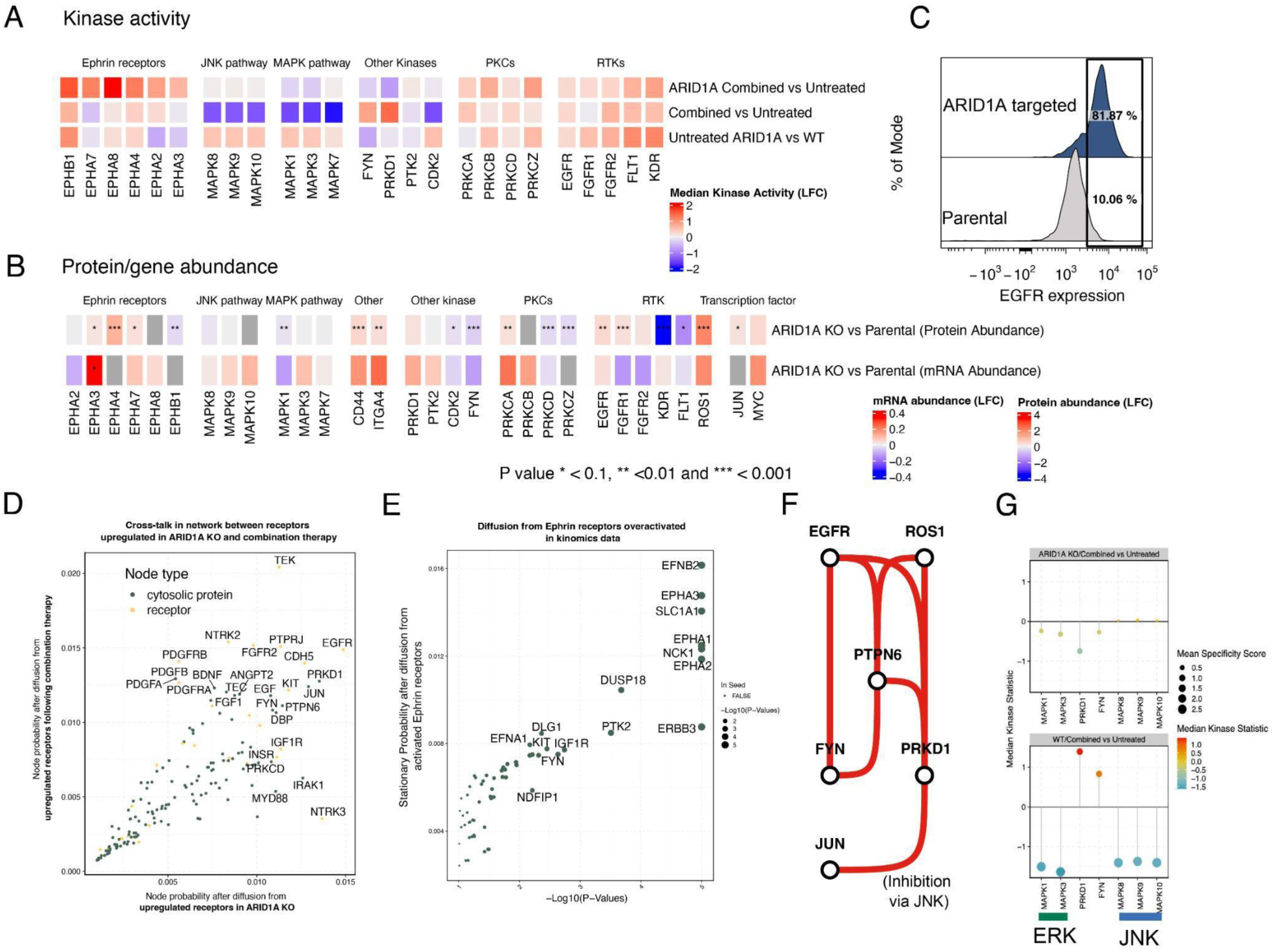
Summary of changes induced by ARID1A KO and their interaction with drug response. **A.** Kinase activity of selected kinases on ARID1A KO and WT at a basal level and post-combination drug treatment. **B.** Heatmap demonstrating the changes in proteomic and transcriptomic abundances for selected kinases at a baseline state for ARID1A targeted compared to parental A375 cell line. **C.** Expression of EGFR as determined by flow cytometry on the surface of ARID1A KO or parental A375 cell line. **D.** Scatter plot showing probability distribution from the network propagation from receptors that are upregulated in response to combination therapy (y axis) and receptors that are upregulated in response to ARID1A KO (x axis). **E.** Scatter plot showing probability distribution from the network propagation from Ephrin receptors activated following treatment of combination therapy in ARID1A KO cells. X axis refers to significance (-log10(P)) and the y axis shows the probability distribution of specific nodes. **F.** Network showing how paths from receptors converge on JUN via FYN and PRKD1 following network propagation from EGFR and ROS1. **G.** Kinase activity assay showing the median kinase activity (y axis) of selected kinases (x axis) following combination drug treatment in parental A375 (ARID1A WT) cell lines (bottom) and ARID1A KO A375 cell lines (top).

To explore the relevance of this transcriptional reprogramming and how it can rewire responses to therapy, we simulated cross-talks from ARID1A KO-induced changes in receptors and therapy-induced receptor changes. We combined networks derived from Factor 1 (representing drug-agnostic changes in signalling) and Factor 3 (representing ARID1A KO induced changes in signalling) into a network that describes the interaction between these two processes (**Fig. S8A**). The resultant network was significantly enriched for the terms ‘MAPK signalling pathway’, ‘Immune system’, ‘Melanoma’, ‘Focal adhesion’ and ‘FGF signalling pathway’ (**Fig. S8B**). To study cross-talks, we performed a Markov Random Walk (*48*) to simulate signal propagation from receptors with increased abundance (**Material and Methods**). This was done separately for those receptors upregulated in ARID1A KO versus receptors increased in combined trametinib and vemurafenib treatment, to identify where the two signals converge, and which proteins are common in both processes. This analysis reveals that the random walk converges on several critical proteins, including the transcription factor JUN, as well as the proteins PRKD1, FYN, PTPN6 and PRKCD (**Fig. 4D**).

We also performed the same random-walk analysis on the members of the Ephrin receptor family that were strongly activated after drug treatment in ARID1A KO cells, but not in the parental cell line (**Fig. 4A and E, and Fig. S5C**). This flagged other receptors, as well as the tyrosine kinase adaptor protein NCK1, the phosphatase DUSP18 and the kinases PTK2, FYN and DAPK. Absolute levels of the mRNA and protein reveal that NCK1 is no longer responsive to drug therapy as it was in the parental cell line, whereas the levels of DUSP18 become responsive to drug therapy in ARID1A KO, but not in the parental cell line. Conversely, the extent of the changes of the abundances of the kinases FYN and DAPK1 do not differ much between genetic conditions. However, their baseline abundance is affected in both cases (**Fig. S8C**).

By calculating maximum flow through the network from these important receptors to MAPK1/3 and JUN/PRKD1, we can assess the routes via which these proteins can interact with each other within our ‘omics’-derived networks (**Fig. S9A and B; Materials and Methods**). A schematic ‘circuit’ showing the simplified pathways of these networks is shown in **Figure 4F**. This shows the intermediary role of PRKD1, FYN and PTPN6 in mediating EGFR/ROS1 signalling towards JUN. According to our functional kinomics data, both kinases FYN and PRKD1 are no longer highly active following drug treatment in ARID1A KO cell lines (**Fig. 4G**).

### ARID1A KO suppresses HLA proteins in both *in vitro* and *in vivo* contexts

Given the role of ARID1A in gene regulation, we then looked specifically at the enrichment of transcriptional regulons in the context of drug treatment in ARID1A KO background cells to identify the differential activity of transcription factors. We see an increase in basal levels of JUN and MYC activities following ARID1A KO, consistent with our previous findings (**Fig. S7C**). Strikingly, in ARID1A KO, we also observe a significant decrease in the expression of the regulons of RFX5, RFXAP and RFXANK, which form the RFX complex to regulate the transcription of MHC class-II genes and NFYC which is also involved in the regulation of the same genes (*49*) (**Fig. 5A**). In ARID1A KO context, we identify a consistent increase in expression of genes that are repressed by RFX5 (COL1A2) and a significant decrease in expression of those genes that are activated by RFX5 (MHC class-II and CD74) (**Fig. 5B**). While MHC class-II family members, which present antigens to CD4^+^ T-helper cells, are typically restricted to professional APCs (antigen-presenting cells), such as dendritic cells and B cells, A375 cells have been shown to express MHC class-II molecules (*50*). To validate our finding, we tested if the expression of MHC class-II proteins on the surface of ARID1A KO A375 cell lines were altered. Remarkably, we found that the expression of HLA-DQ and HLA-DR was significantly decreased in ARID1A KO cells (**Fig. 5C**). Of note, among the upregulated transcription factor (TF) activities in ARID1A cells, we were able to recognize TFs in mediating the regulation of extracellular matrices, with TWISTs and SMADs being characteristic examples (**Fig. 5A and B**).

**Figure 5.**
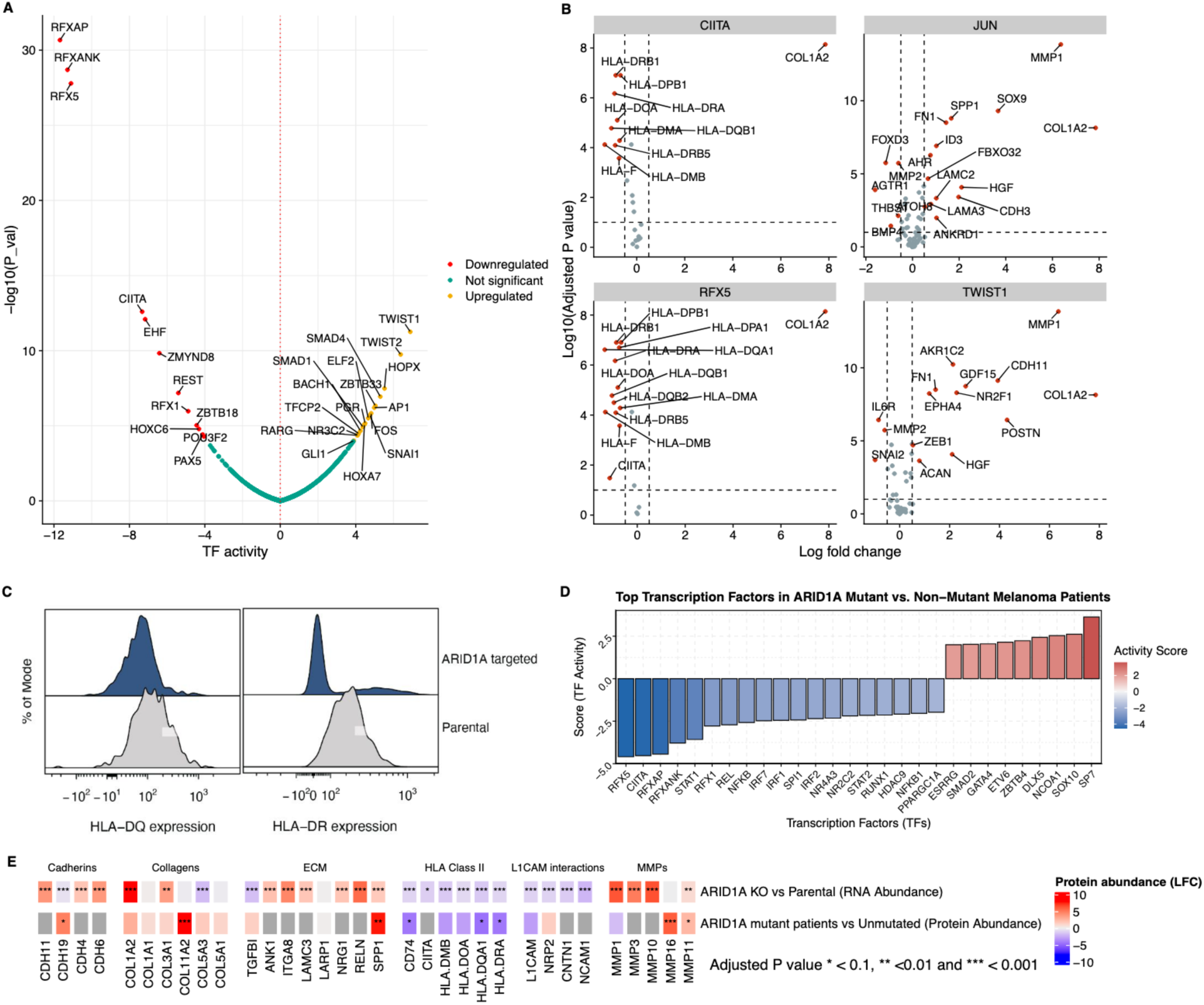
Overview of molecular signature associated specifically with combination drug response (Factor 2). **A.** Transcription factor activity changes (x axis) in ARID1A-targeted A375 cell line compared to parental line. **B.** Log-fold changes for regulons (x axis) of four key transcriptional regulators (CIITA, JUN, RFX5, and TWIST1) that were identified to have significantly differential (y axis) activity at a basal state for the ARID1A KO cell line compared to the parental (ARID1A WT) A375 cell line. **C.** Cell surface expression of two MHC class-II antigens (HLA-DQ and HLA-DR) on parental and ARID1A KO cell lines. **D.** Top transcription factors and regulators (e.g., CIITA) with differential activity (y axis) in ARID1A-mutant versus non-mutant melanoma patients from TCGA. **E.** Heatmap demonstrating the changes in proteomic and transcriptomic abundances (rows) for selected genes/proteins (columns) at a baseline state for ARID1A-targeted (KO) compared to parental (WT) A375 cell line.

Using the TCGA database (https://www.cancer.gov/tcga), we explored whether these same patterns could be identified in patient data. We stratified 472 melanoma patients into ARID1A affected and unaffected groups by finding patients with either mutations or deletions in the *ARID1A* gene (see **Materials and Methods**). This identified 80 patients with genetic aberrations in *ARID1A* and 392 patients without such aberrations (**Fig. S10**). By studying the gene expression of members of the regulons of transcription factors, we found that these patients have significantly lower predicted activities of the MHC class-II regulators RFXAP, RFX5 and CIITA (adjusted *p* value < 0.001) (**Fig. 5D**). Furthermore, we find a significantly elevated activity of the transcription factor SP7, a protein regulating the expression of collagens and metalloproteins. Studying individual gene expression, we reveal that both ARID1A-affected patients and ARID1A KO cells have increased expression of collagens and laminins, which can contribute to increased ECM stiffness (*51*), as well as a decrease in HLA proteins (**Fig. 5E**).

## Discussion

Resistance to MAPK inhibitors poses a major challenge in melanoma treatment, driven by adaptive and acquired mechanisms that enable tumour cells to evade therapy. While prior studies often focus on either adaptive responses or acquired resistance in narrow contexts, we examined early signalling changes after drug treatment and compared them to responses in melanoma cells with acquired resistance. Using *ARID1A*, a resistance-associated gene frequently mutated in melanoma, we explored short-term cellular responses in distinct genetic contexts. ARID1A’s role in resistance and immune evasion underscores its therapeutic relevance, making it a compelling focus for this study.

In our study, we adopted an integrative approach to uncover network-level insights into how signalling responses change under drug pressure in cells with two distinct genetic states. By analyzing transcriptomics, proteomics, and signalling datasets, we addressed the limitations of single-metric approaches. Transcriptomics data offers broader coverage, while proteomics and phosphoproteomics are more affected by technical and biological noise (*52*). To integrate these datasets effectively, we combined integrative matrix factorization (*33*) with a network-centric method that we had previously developed (*34*), enabling robust interpretation of complex data. This approach highlighted differences in signalling pathways influenced by drug treatment and genetic alterations.

We focused our study on early cellular responses to drug treatment, selecting the 6-hour time point as it maintains the pathway inhibition intact and allows the observation of critical cellular changes, such as reduction in the abundance of dual-specificity phosphatases (DUSPs) and other MAPK cascade negative regulators (*20*). This loss in phosphatases increases cell signalling activity by impairing signal termination from RTKs. We confirmed this feedback sensitivity by observing downregulation of *DUSP*, *SPRY* and *SPRED* genes within 6 hours of drug treatment, alongside heightened RTK (EGFR, IGFR, FGFR) kinase activity, despite minimal changes in their gene or protein expression levels.

We used our functional kinomics dataset as an independent and orthogonal way to validate signalling pathways inferred from our multi-omics integration approach. We found that the key nodes in our factor 1-derived network were also the ones whose activity changed the most (mainly for serine/threonine kinases), with a notable decrease in MAPK1/3 activity (the drug target). One striking change both at gene expression level and kinase activity level was that for PRKD1. PRKD1 is a member of the PKD serine/threonine kinase family that can be activated downstream of RTKs and, also, in response to increase in cellular reactive oxygen species (ROS). It has been long known to be a suppressor of epidermal growth factor (EGF)-dependent JNK activation (*53*) by directly complexing with JNK (*54*). Upon PKD phosphorylation by PKC, it complexes with JNK and inhibits its ability to phosphorylate c-Jun at a critical serine-63 position. PRKD1 is frequently mutated and highly expressed in melanoma relative to other cancers, with pro-proliferative or anti-proliferative effects being dependent on context (*55*). We noted an increase in both expression and activity of PRKD1 upon treatment with drugs. This was consistent with decrease in JNK phosphorylation indicating decreased JNK/c-Jun axis activity. In mutant-BRAF melanoma, the JNK/c-Jun signalling pathway is associated with apoptosis (*19*) and pathway activity increases in cells that are quiescent and resistant to apoptosis.

Combination therapy is used in clinics targeting two separate entities in the dependent MAPK pathway and has been shown to provide longer efficacy. A paradoxical activation of the MAPK pathway in BRAF non-mutant cells when BRAF alone is targeted has been described before, which further justifies targeting both BRAF and downstream MAPK components (MAP2K1/2) (*56*). At a signalling level, we detected small but specific changes in the phosphorylation of proteins related to DNA repair pathways, such as RIF1, and to EMT, such as SIRT1 and LEO1, although we could not account for these changes to phenotypic variation, as in our experimental design with BRAF-mutant melanoma, we did not observe a significant difference in killing rates with mono or combination therapies. Given our observations, more research is needed to establish whether such changes have indeed a role in promoting EMT and affecting immunogenicity of melanoma cells upon combination treatment.

We noted that ARID1A KO and parental cells showed similar mRNA and protein expression responses to drug treatment, including intact feedback mechanisms like DUSP downregulation. However, signalling outcomes differed: ARID1A KO cells maintained MAPK1/3 and JNK activity, indicating resistance to MAPK pathway inhibition. In parental A375 cells, PRKD1 was activated by drug treatment, while in ARID1A KO cells, its activation was suppressed, likely due to transcriptional rewiring. We propose that this relieved JNK inhibition, leading to increased JUN activity, a key node in the resistant network. JUN upregulation is a common response in BRAF inhibitor-treated melanomas (*19*), and dual targeting of JUN and BRAF has shown synergy in overcoming resistance (*57*).

In our multi-omics network analysis, we identified two different types of signalling behaviour at the receptor level in ARID1A KO cells, which could contribute to this inability to respond to MAPK pathway inhibition. This included differences at a basal expression level of RTKs (EGFR, ROS1) but also at a signalling effect, post drug perturbation level, specifically driven by Ephrin receptors. Studies have demonstrated that elevated expression of receptor tyrosine kinases (RTKs), such as EGFR, can overwhelm the mechanisms responsible for receptor endocytosis and degradation. As a result, the receptor remains chronically active rather than exhibiting transient activation in the presence of a ligand (*58*, *59*). Recent work using mechanistic modelling shows that when oncogenic BRAF is fully inhibited, MAPK pathways can be turned on in a RAS dependent manner, which, in turn, is stimulated by receptor tyrosine kinase (RTKs) (*14*). We propose that the basal increase in RTKs (EGFR, ROS1) in ARID1A KO cells and the increased activity from Ephrin receptors post drug treatment could monopolise cellular signalling, leading to changes in PKC dynamics and attenuate PRKD1 response upon treatment with MAPK inhibitors, which leads to higher baseline JUN activity. This is consistent with our observation that subunits of PKC (PRKCB and PRKCZ) themselves are already at a higher phosphorylation status in the ARID1A KO cell line.

How mutations in ARID1A influence tumour-immune interactions has been a field of study for multiple cancer types. Previous studies have shown that ARID1A mutations disrupt interferon (IFN) signalling, which diminishes cytotoxic T-cell infiltration, leading to compromised effectiveness of immunotherapy models (*30*). Notably, our analysis of ARID1A-targeted cells and patients with ARID1A mutations revealed significant deficiencies in the basal expression of HLA-related proteins, which are regulated by IFN signalling (*60*). Apart from loss in MHC class II expression, we also observed enrichment of ECM component expression in these mutant cells. Intriguingly, the same classes of proteins (collagens, laminins, MMPs) have been previously reported to be differentially upregulated in anti-PD1 treatment resistant MC-38 cell lines in mice compared to their treatment-sensitive counterparts (*61*, *62*), suggesting that this basal signature of ARID1A could reduce the efficacy of T cell infiltration and yield immunotherapy less efficacious.

Like previous studies (*36*), we observe the key importance of receptors in rewiring downstream signalling towards resistance. However, in our data, we detect no evidence that ARID1A perturbs any of the negative feedback mechanisms by Gerosa *et al.* (2020) (*36*). However, in our system-level characterisation, we see widespread increases in multiple receptors, including EGFR, ROS1, FGFR1 and ITGA4. This suggests a mechanistic redundancy that makes selecting a single ‘silver-bullet’ protein to target infeasible as cellular signalling has multiple routes to restore lost signalling. We used network-centric methods to propagate from these receptors within our network, to reveal new proteins with uncharacterised associations to MAPK resistance, including FYN, PRKD1 and NCK1. NCK1 is an adaptor protein closely associated with Ephrin signalling (through which it was flagged in this analysis). We find that its abundance is distinctly affected by drug response in the parental A375 cell lines, versus ARID1A KO cells. In WT A375 cells, its (*NCK1*) gene abundance drastically drops following drug treatment, whereas when ARID1A is knocked out, it remains unresponsive. In non-oncogenic contexts, NCK1 is a known activator of both JUN (through JNK (*63*)) and MAPK (through RAS (*64*)). In this data, we find both these families of kinases to be unresponsive upon ARID1A KO, suggesting a functional interaction between these two events consistent with prior literature.

In this study, we used innovative graph-theoretical techniques to integrate multi-omics data using networks. Prior research has arrived on a similar method, showing the utility and potential of using MOFA to construct networks from integrated multi-omics datasets (*65*). By combining factor analysis with diffusion-based network construction, our study uniquely incorporates phosphoproteomics, transcriptomics and protein abundance data to derive unbiased, multi-omics networks that reveal perturbed pathways in melanoma drug resistance. Unlike methods relying solely on annotated pathways, this approach allows the data to drive the discovery process, while still enabling functional annotation through enrichment analysis. Additionally, the inclusion of kinomics activity data validates the identified network nodes by linking them to the most perturbed kinases following drug treatment. Our methodology captures the orthogonal effects of drug treatments and genetic rewiring, while uncovering shared protein-protein interactions that describe the interplay between the two processes, offering a robust framework for extracting biologically relevant insights from noisy datasets and prioritising hypotheses for further investigation.

In summary, this study provides a comprehensive systems-level analysis of the early cellular responses to MAPK pathway inhibition in melanoma, revealing distinct adaptive and resistance mechanisms. By leveraging innovative graph-theoretical and multi-omics integration techniques, we demonstrated that ARID1A-mediated transcriptional rewiring significantly alters the cellular signalling landscape, driving resistance to MAPK inhibitors. Our findings highlight key differences between parental and ARID1A KO melanoma cells, particularly in signalling pathways involving PRKD1, JUN and receptor tyrosine kinases (e.g., EGFR, ROS1). These differences underpin the enhanced resistance observed in ARID1A KO cells. Additionally, the integration of functional kinomics with multi-omics datasets enabled the identification of critical nodes within drug-resistant signalling networks. Notably, we observed a mechanistic redundancy in receptor activation that complicates single-target therapies but presents opportunities for combination strategies targeting JUN or immune pathways. These insights advance our understanding of melanoma resistance mechanisms and lay the groundwork for more effective therapeutic interventions, including the potential integration of immunotherapies tailored to specific genetic and signalling contexts.

## Acknowledgements

The work was supported by the European Molecular Biology Laboratory (C.G.B., S.S, E.P) and the EMBL International PhD Programme (C.G.B.); The EMBL GeneCore Facility is acknowledged for support in transcriptomics data collection. EMBL IT Support is acknowledged for provision of computer and data storage servers. The proteomics and phosphoproteomics data collection was supported by the EU Horizon 2020 program INFRAIA project Epic-XS (Project 823839). This research was also funded by the Wellcome Trust (grant 206194) to G.J.W.

## Author contributions

**CGB:** Conceptualization, Methodology, Software, Formal analysis, Investigation,Visualization, Writing - Original Draft, Writing - Review & Editing; **SS:** Conceptualization, Validation, Methodology, Formal analysis, Investigation,Visualization, Funding acquisition, Writing - Original Draft, Writing - Review & Editing; **AMS:** Validation, Investigation; **KSN:** Investigation; **ADV:** Investigation, Formal analysis; **FIV:**Investigation, **AM:**Validation, **VP:**Validation, **MA:** Resources, Supervision **GJW:** Resources, **LGA:**Validation,Resources **SJD:** Resources, **DJS:** Resources, Supervision, Writing - Review & Editing **EP:** Conceptualization, Methodology, Resources, Funding acquisition, Project administration, Writing - Original Draft,Writing - Review & Editing

## Materials and Methods

### Cell culture

Parental and *ARID1A*-targeted A375 lines were grown in DMEM-F12 (Gibco, Cat. No: 11320033) with 10% FBS (Gibco, Cat. No: 10500-064) and 1% Penicillin-Streptomycin-Neomycin (Sigma, Cat. No: P4083) at 37°C with 5% CO2. Cells were grown in a monolayer, and the culture medium was changed every 3 d. The cells were passaged once they reached ∼80% confluence.

### Whole genome CRISPR screen in the presence of trametinib

A genome-wide screen to identify genes conferring resistance to trametinib was performed using a human genome-wide library (Yusa V1), which targets ∼18,000 genes with ∼91,000 gRNAs using a detailed protocol described with reagents and product codes previously (*66*). In short, two sets of Cas9 expressing A375 cells (80 million starting population for each set) were transduced at a MOI of 0.3 with the genome-wide lentiviral library. A day post infection, cells were treated with 2 μg/mL puromycin to remove any non-transduced cells. Live cells on day 7 were split into 2 populations; one was treated with 1 nM trametinib and the control set was left untreated. All live cells post-treatment, and 30 million control untreated cells, were collected after 2 weeks, genomic DNA was extracted, gRNAs were amplified and libraries were generated. The MAGeCK software (*67*) was used to identify genes that were enriched in the live population compared to the control population.

### Luminex assay

For the Luminex assay, roughly 20,000 cells were seeded on 96-well plates overnight in 100 μL serum-starved medium (Cell culture media without FBS). In the next morning, culture media containing serum (standard cell culture media) and supplemented with drugs or growth factors (as relevant) were added to the cells. After 6 hours, the plate with cells was placed on ice and cells washed with 100 μL of cold PBS. Next, 60 μL lysis mix was added to the cells and the plate was shaken for 20 minutes at +4 °C and 650 rpm. The lysis mix was prepared by mixing Lysis buffer (Protavio, Cat. No: PR-ASSB), Protease Inhibitors (1 tablet to 50 mL of Lysis buffer, Roche, Cat. No: 11873580001) and 2 mM PMSF (Sigma, Cat. No: 93482). Cell lysates were then frozen at -20 °C and phosphorylation was measured utilising the multiplex assay service provided by Protavio (*68*) (Athens, Greece). We developed a phospho-plex platform for semi-quantitative analysis of the phosphorylation status of 17 phosphoproteins (**Table S7**), which displays a good signal to noise ratio to be measured in the *in vitro* assays (**Table S8**).

The assay is based on the xMAP technology developed by Luminex Corporation. A mix of 17 capture antibodies coupled to Luminex magnetic beads (Bead mix) was prepared. Each antibody is coupled to a different magnetic bead region. Beads can be uniquely identified and differentiated by the Luminex instrumentation due to their unique color classification. A ‘detection’ mix consisting of 17 biotinylated secondary antibodies specific for recognizing the analytes of the panel was also prepared. Detection antibodies are biotinylated, in order to be recognized by a streptavidin-phycoerythrin (SAPE) substrate used to produce the final detection signal.

Each sample is incubated with the bead mix in a well of a 96-well microtiter plate to allow binding of the analyte. Any unbound material is removed by washing using a magnetic separator. The formed antibody-analyte complex is incubated with the secondary detection antibody mix. Any unbound detection antibody is removed by a washing step and the formed complex of capture antibody-analyte-detection antibody is labelled with SAPE (MOSS Inc., Pasadena, Maryland, USA, Cat. No: SAPE-001). The fluorescent emission of R-phycoerythrin and the distinct microsphere fluorescent signatures are measured simultaneously by the Luminex^®^ instrument.

### Generation of ARID1A KO cell line

ARID1A-targeted cell lines were purchased from Synthego. Sequencing across the cut locus revealed a single nucleotide insertion, which led to a frame-shift mutation (**Fig. S1A**). The KO efficiency as determined by TIDE was ∼98% (**Fig. S1B**) (*69*).

### Sample preparation for phosphoproteomics and functional kinomics analysis

Cells were seeded overnight at 0.5 x 10^6^ cells/well density in 6-well plates. Before harvesting, cells were treated with vemurafenib (APExBio, Cat. No: A3004) (1 mM) and trametinib (APExBIO, Cat. No: A3018) (10 nM), separately or in combination, for 6 hours. Approximately 10^6^ cells were used as the starting material. Cells were washed twice with ice-cold PBS, scrapped with a cell scraper, and then centrifuged at 1000 × *g* for 3 min. The supernatant was removed, and the cell pellets were frozen in liquid nitrogen. For functional kinomics data generation, the same protocol was used except that 2 x 10^6^ cells/T25 flasks were seeded overnight and all cells were collected post-treatment.

### Kinome activity profiling through functional kinomics

Protein Tyrosine Kinase (PTK) and Serine/Threonine Kinase (STK) activity profiles were assessed using PamChip® peptide microarrays (Pamgene International BV, BJ’s-Hertogenbosch, The Netherlands), which measure the ability of active kinases in a protein lysate to phosphorylate specific peptides imprinted on multiple peptide arrays. A typical PTK PamChip® microarray contains 196 immobilized peptides, while a STK PamChip® microarray contains 140 peptides covalently attached to a porous material. Peptides harbor phosphorylation sites derived from literature or computational predictions that are associated with one or more upstream kinases. Phosphorylation is detected by (phospho-)specific primary antibodies and the signal is quantified by FITC-conjugated secondary antibodies. Detection is performed in multiple cycles at different exposure times and is monitored by a CCD camera. Protein lysates were prepared from untreated and drug-treated A375 melanoma cell lines of both genotypes, using M-PER mammalian extraction buffer (Thermo Fisher Scientific, Waltham, MA, USA), containing Phosphatase Inhibitor Cocktail and Halt Protease Inhibitor Cocktail EDTA-free (1:100 each; Thermo Fisher Scientific, Waltham, MA, USA), and quantified by the Bradford assay. A total of 5 μg and 1 μg of protein lysates per array from each biological sample were used to profile tyrosine and serine/threonine kinase activity, respectively, according to PamGene’s standard protocols. Image analysis, peptide quality control (QC), signal quantitation, data normalization and visualization, as well as upstream kinase prediction were performed using Bionavigator v.6.3 software (PamGene, ‘s-Hertogenbosch, The Netherlands), according to manufacturer’s instructions. Three independent biological replicates were used for each condition (parental (wild-type, WT), ARID1A KO, untreated or drug-treated A375 melanoma cells).

### Transcriptomics data generation

Cells were seeded at 0.5 x 10^6^ cells/well density in 6-well plates and left overnight. Before harvesting, cells were treated with Vemurafenib and Trametinib, separately or in combination, for 6 hours. Approximately 10^6^ cells were used as the starting material. Cells were washed twice with ice-cold PBS, scrapped with a cell scraper, and then centrifuged at 1000 × *g* for 3 min. For RNA isolation, the RNeasy Mini Kit (Qiagen, Cat. No. 74104) was used per manufacturer instructions, along with QIAshredder columns for the homogenisation of cell lysates and DNase I treatment using DNase I kit (Thermo Fisher, Cat. No. ENO525). Following RNA extraction, the integrity of the isolated RNA was assessed using the Agilent 4200 TapeStation system (Agilent) according to manufacturer’s instructions, using 2 μL of a 1:150 dilution in H2O of each purified total RNA sample. Samples were tested further for cDNA synthesis using PrimeScript 1st strand cDNA synthesis kit (Takara Bio, Cat. No. 6110A) followed by RT-PCR on *β*-Actin using 2 μL of cDNA product.

### Transcriptomics data processing

We performed normalisation of the raw counts using the voom function from the limma package (*70*). We selected only genes with a count-per-million (CPM) greater than 2 in at least one sample, to retain for downstream analysis. We computed scaling factors to convert observed library sizes into effective library sizes using the library edgeR (*71*). We used these normalised counts as inputs to MOFA (*33*). For visualisation purposes we used the package limma to predict stable results and to calculate significance provided for the presented volcano plots and LFCs (Fig. S3A and Tables S1-3).

For transcription factor activity inference, we used the Python (version 3.8.18) package decoupleR. Differential expression was computed using the Wald test, and log-fold changes (LFCs) were extracted using the pydeseq2 library. The CollecTRI gene regulatory network, focusing on human transcriptional regulation, was retrieved using the decoupler library. We employed the Univariate Linear Model (ULM) implemented in decoupler to infer transcription factor (TF) activities from differential expression data (*72*).

### Proteomics/phosphoproteomics data generation

#### Sample preparation

300 μL of a detergent-based buffer (1% sodium deoxycholate (SDC), 10 mM tris(2-carboxyethyl)phosphine (TCEP), 10 mM Tris, and 40 mM chloroacetamide) with cOmplete mini EDTA-free protease inhibitor cocktail (Roche Cat. No. 04693132001) was added to the cell lysates, boiled for 5 min at 95 °C, and sonicated using the Bioruptor for 20 cycles of 30 sec on : 30 sec off. Protein quantification was carried out using the Bradford assay and an aliquot corresponding to 200 ug was retained for each sample. 50 mM ammonium bicarbonate was added, and digestion was allowed to proceed overnight at 37 °C using trypsin (Sigma **T6567**) and LysC (Wako, 125-05061) at 1 : 50 and 1 : 75 enzyme : substrate ratios, respectively. The digestion was quenched with 10% formic acid and the resulting peptides were cleaned-up in an automated fashion using the AssayMap Bravo platform (Agilent Technologies) with a corresponding AssayMap C18 reverse-phase column, followed by vacuum drying. To generate a reference channel to be used for all experiments, a pool of all samples combined was digested.

Dried peptides were re-solubilised in TMT resuspension buffer (87.5% HEPES, 12.5% ACN) and 11-plex TMT labels (Thermo Fisher A34808) were prepared according to manufacturer’s instructions. TMT labels were added to samples and labelling was allowed to occur during 2 hours at RT (room temperature), after which the reaction was quenched using 5% hydroxylamine in HEPES, for 15 minutes at RT. The various channels were combined for each experiment and the acetonitrile content was reduced by evaporation. The samples were then cleaned using a SepPak 1cc cartridge and dried completely before solubilising in HpH buffer A (10 mM NH4OH, pH 10.8). HpH fractions were collected every minute over a 100 min gradient. Fractions between minutes 10 and 70 were concatenated into 20 fractions, and an aliquot of each was set aside for vacuum drying and full proteome analysis. The remainder of the concatenated fractions were vacuum dried followed by re-solubilization in 80% ACN/ 0.1% TFA for phosphopeptide enrichment. The enrichment was carried out in an automated fashion using the AssayMap Bravo platform (Agilent Technologies) with corresponding AssayMap Fe(III)-NTA cartridges, and eluates were dried by vacuum centrifugation and resolubilised in 1% FA, of which ∼1 μg was injected on column.

#### MS analysis

All spectra were acquired on an Orbitrap Exploris mass spectrometer (Thermo Fisher Scientific BRE725535) coupled to an Ultimate 3000 liquid chromatography system. Peptides were trapped on a 300 µm i.d. x 5mm C18 PepMap 100, 5 µm, 100 Å trap column (Thermo Scientific P/N **160434**) and then separated on a 50 cm (75 um ID) in-house packed column using Poroshell 120 EC-C18 2.7-Micron (ZORBAX Chromatographic Packing, Agilent). Samples were eluted over a linear gradient ranging from 9 - 45% 80% ACN / 0.1% FA over 65 min, 45 - 99% 80% ACN / 0.1% FA for 3 min, followed by maintaining at 99% 80% ACN / 0.1% FA for 5 min at a flow rate of 300 nL/min. Phosphopeptide-enriched samples were eluted over a linear gradient ranging from 9 - 34% 80% ACN / 0.1% FA over 95 min, 45 - 99% 80% ACN / 0.1% FA for 3 min, followed by maintaining at 99% 80% ACN / 0.1% FA for 5 min at a flow rate of 300 nL/min. MS1 scans were carried out at a resolution of 60000, with standard AGC target and automatic IT. The intensity threshold was set to 5.0e4, charge states 2-6 were included, and dynamic exclusion was used for a duration of 14 sec. For the MS2 scans, a window of 1.2 m/z was applied, with HCD collision energy set to 30%, an orbitrap resolution of 45000, and standard AGC target and automatic maximum injection time determination.

#### Raw data processing

All raw files were processed using ProteomeDiscoverer (Thermo) version 2.4.1.15 and obtained data were searched against the SwissProt Homo Sapiens proteome (April 2021 release), using the Mascot proteomic search engine with the following settings: a maximum of 2 missed cleavages, precursor mass tolerance of 10 ppm and fragment mass tolerance of 0.8 Da. Oxidation (M) and Phosphorylation (STY) were selected as dynamic modifications with carbamidomethyl (C)- and TMT-tags (K and N-terminal) being selected as static modifications.

### Phosphoproteomics data processing

Data processing and analysis were conducted in R (version 4.2.0) (*73*). We took raw peptide abundances and protein abundances from Proteome Explorer, and we normalised both by sample loading normalisation. The normalised data was processed to extract phosphosite information for peptides, including amino acid residue and position. UniProt IDs and phosphosite information were parsed using the EnsDb.Hsapiens.v86 annotation R package (*74*).

The data is sparse, so missing values were imputed across replicates within a single condition using the function scImpute from the R library PhosR (*75*). We performed this if there are 3 or more quantified values of that variable within a given condition (drug treatments and ARID1A KO). Data was median centred and scaled, and batch effects from the different TMT-11 plex runs were handled by the R package ComBat (*76*). Only samples with quantified values above a threshold of 50 were retained for further analysis. Log2 transformations were applied to facilitate downstream statistical modelling.

To decouple the effects of total protein abundance on phosphorylation, a regression-based method was employed to estimate the “net” phosphorylation level (as implemented in (*77*)). This approach calculates residuals from a linear model where log-transformed phosphoprotein abundance is regressed against the corresponding total protein abundance for each sample (*phosphopeptide abundance ∼ protein abundance*). The resulting residuals represent phosphorylation changes independent of total protein levels. The processed data was aggregated based on unique phosphosite identifiers, with any duplicate entries being combined.

### Multi-omics factor analysis

To input the multi-omics data into Multi Omics Factor Analysis (MOFA) (*33*), we used ANOVA to select for the most highly variable phosphopeptides and proteins. The criteria for phosphopeptides significance were an FDR adjusted ANOVA *p*-value < 0.1 and an absolute mean abundance change > 0.5. These were then standardised by z-score transformation. The criteria for protein and mRNA abundance were more strict (those with *p*-values < 0.001 and absolute values > 1). This helps in reducing noise by focusing on biologically relevant features.

A MOFA object was created using the integrated, long-format dataset, comprising phosphoproteomic, proteomic, and mRNA expression views. Default data, model and training options were configured using the R package MOFA2 (*78*). The number of latent factors was set to 3, reflecting the number of sources of variation in the data (**Fig. 1** and **Fig. S4**). After training, sample metadata (e.g., drug treatment and genetic background) were integrated into the MOFA object, enabling stratified analysis based on biological conditions. The feature loadings for Factor 1 were multiplied by -1, such that in all factors negative loadings represent features that correspond to control samples (untreated, non-combination therapy or WT in factors 1, 2 and 3, respectively).

### Network Propagation using phuEGO

To explore the functional interactions between proteins identified from the MOFA factors, we employed phuEGO, a tool designed for signalling network construction (*34*). The goal was to identify potential regulatory hubs and pathways that are influenced by the key proteins associated with the latent factors across all omics views in our dataset. Latent factor weights were extracted for each omics view (i.e., phosphoproteomics, proteomics and mRNA expression) using the get-weights function from the MOFA2 package. For each protein, the phosphosite with the highest absolute weight was selected and then the results of each condition were aggregated across each view, so we had a protein-level weight. We extracted proteins exhibiting the most extreme weights for each factor by selecting the top or bottom 5% quantiles. Using these proteins as seeds, we ran phuEGO with parameters Fisher-threshold = 0.1, Fisher-background = intact, Random-walk damping = 0.95, RWR-threshold = 0.01 and KDE-cutoff = [0.5]. This generated a network for each factor found in the multi-omics data.

### Prize collecting Steiner-Forest analyses

To identify key functional genes within the larger factor networks, we employed the Prize-Collecting Steiner-Forest (PCSF) algorithm with randomised edge costs. The randomised method enhances the robustness of the resulting sub-network by running the algorithm multiple times, while adding random noise to the edge costs (*79*). We selected the top 50 central genes as terminal nodes based on their PageRank centrality, which has been proven to produce biologically meaningful results (*80–83*). These nodes were then weighted as prizes and then the PCSF algorithm was run with 4,000 iterations, with up to 5% random noise added to edge costs.

### Random walk of receptors upregulated in ARID1A KO cancer cells

We implemented an algorithm for analysing receptor-mediated pathways using network propagation (via random walks), with a focus on perturbations caused by ARID1A-knockout (KO) and combination drug treatments. We used the union of graphs for Factor 1 (showing drug agnostic responses) and Factor 3 (showing ARID1A KO-induced responses) to visualise the interaction between these two processes. For both ARID1A KO and combination drug treatments, we extract receptor LFC (log-fold change) data and split them into upregulated and downregulated receptors. We processed LFC datasets from differential expression analysis by filtering receptors (as defined by Omnipath) that are significantly affected (adjusted *p*-value < 0.01). Receptors were identified using the OmnipathR package. These LFCs are normalised by dividing each element by the sum of the vector (to prepare for random walks). We then perform a random walk on a graph with a given starting vector of probabilities derived from the LFCs above. The random walk is corrected for hubs and used to compute a stationary distribution that indicates the relative importance of a node for each condition.

To study the effect of increases in Ephrin receptor activation on our network, we take the union network prior to PageRank/PCSF pre-processing and perform random walk with restart on the receptor kinase activity, determined by the functional kinomics, target-peptide, phospho-tyrosine data. We identified receptor proteins in the kinomics phospho-tyrosine data using the OmnipathR package. We filtered this data to retain kinases that had statistically significant changes in activity (e.g., higher than a threshold of ±1 median kinase activity). Starting from an initial set of deregulated receptors (normalised from the previous step), we performed random walks over the network to estimate the stationary distribution of each node. To assess the significance of observed distributions, we conducted permutation tests by generating 10,000 random networks that preserved the degree distribution of the original network. We compared the original and randomised distributions to calculate *p*-values.

### Maximum flow of receptors to nodes of interest

We selected the nodes that were the most prominent in the heat diffusion from both combination therapy and ARID1A KO (cytosolic proteins, JUN, PRKD1, PTPN6 and FYN) and performed heat diffusion individually from each upregulated receptors in both conditions to detect which receptors were responsible for their flagging (**Fig. S9A**). To identify pathways in the network that are responsible for the highlighted network propagations between these receptors (ROS1, FGFR1 and EGFR) and their targets, we computed the maximum flow between a specified source and sink node from the combined graph, where edge capacities were defined by semantic similarity. This was done using the R package, igraph (*84*).

We extracted the flow values from the maximum flow result and identified edges with flow values exceeding the 95^th^ percentile. For visualisation, a sub-graph was induced from the original graph, retaining only the nodes connected by high-flow edges. We identified the connected components in the sub-graph and selected the component containing both the source and sink nodes. This sub-graph represents the largest connected high-flow region within the network between modulated receptors and nodes of interest.

### Clinical analysis of *ARID1A* gene-perturbed patients

Melanoma RNA-seq data and corresponding mutation information were retrieved from TCGA (The Cancer Genome Atlas) (*85*) using the TCGAretriever package. We extracted mRNA expression data (FPKM values) for melanoma patients using the fetch_all_tcgadata() function. We obtained *ARID1A* mutation and CNV (copy number variation) status to identify samples with *ARID1A* alterations. We then categorised samples as having *ARID1A* mutations, deletions, or neither (wild-type, WT). To determine transcription factor activity, we employed the decoupleR package, using the Univariate Linear Model (ULM) as described above.

We also performed differential expression analysis to identify dysregulated genes in *ARID1A*-mutant versus WT melanoma patients using the limma package. We filtered out low-expression genes (minimum count = 10) using the filterByExpr using the edgeR package. We log-transformed the data and normalised them via engagement of the voom function to stabilise the variance. We then fitted a linear model to identify the differentially expressed genes and identified significant genes on the adjusted *p*-values (FDR < 0.01).

### Enrichment analyses

EnrichR (available https://maayanlab.cloud/Enrichr/) was used to perform enrichment analysis of the members of the network using the BioPlanet 2019 database.

### Data visualisation

All analyses were performed in R or Python, with the code being available upon request. The MOFA analysis was conducted using the MOFA2 package, with additional processing and visualisation supported by dplyr, purrr, stringr, ggplot2, PhosR and EnsDb.Hsapiens.v86.

The ggraph package was used to visualise the receptor networks and the results of the network propagation analysis (*86*).

