## Supplementary Figures for "ARID1A-induced transcriptional reprogramming rewires signalling responses to drug treatment in melanoma"

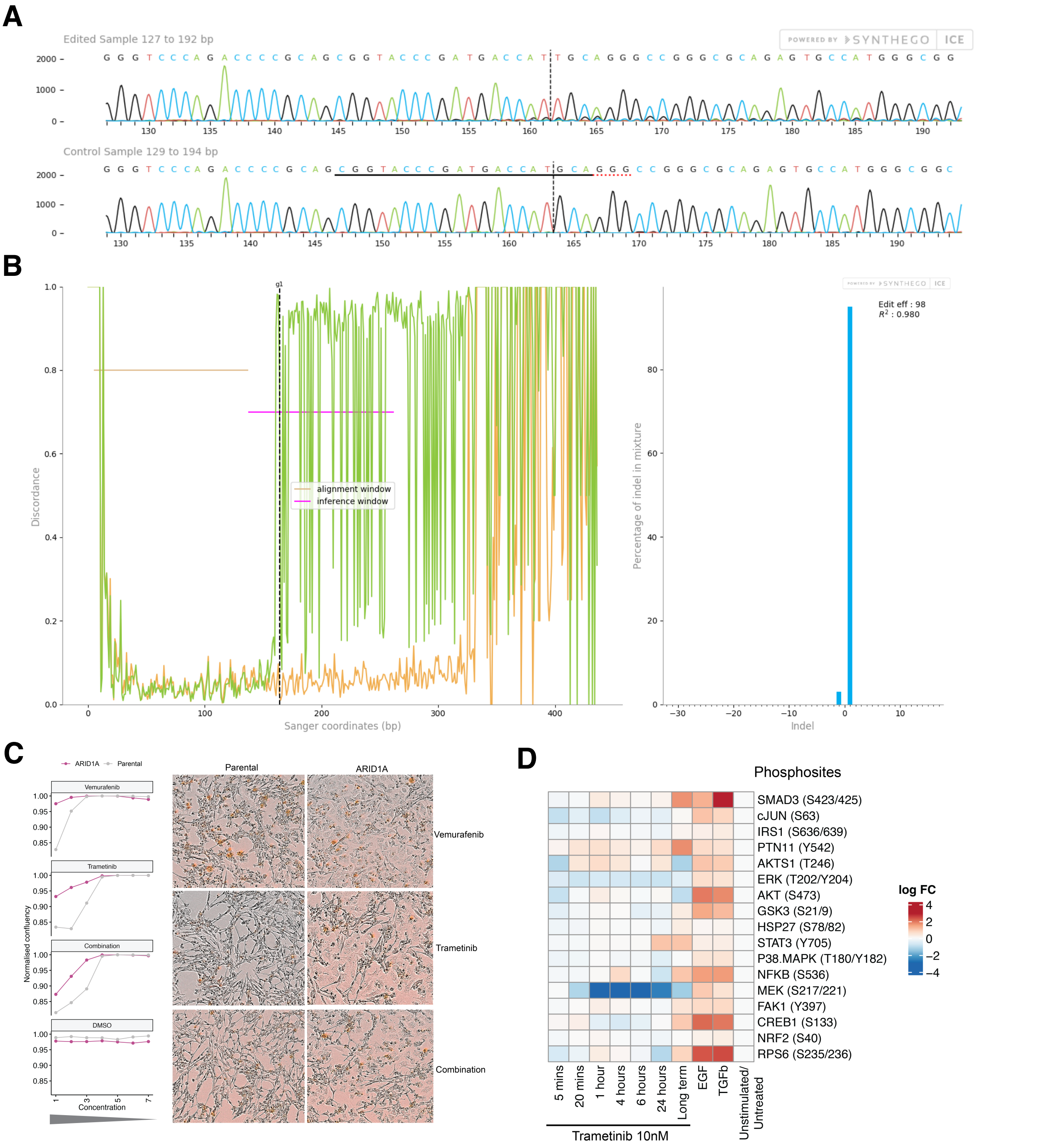


**Supplementary Figure S1. ARID1A was effectively knocked out and affects signalling dynamics. A.** ARID1A and matched parental melanoma A375 cells were purchased from Synthego. Sanger sequencing traces for the edited region is shown. The single base-pair deletion led to a frame-shift mutation in the protein. **B.** Output from TIDE analysis showing mismatches in the inference window (left panel) and 98% editing efficiency. **C.** Killing of parental and ARID1A cell lines in the presence of Vermurafenib, Trametinib, combination of both drugs, and control with DMSO. Starting concentration of 30 nM Vermurafenib and 3 nM Trametinib was used (Left panel). The numbers on x-axis indicate the dilution factor (1:3) from the starting concentration. Images of cells at 90 hours post treatment with the drugs(Right panel). **D.** Luminex-based phosphoprotein measurements of indicated phosphosites on A375 cells treated either with Trametinib (10 nM) for indicated times, or with growth factors (TGFb and EGF) for 4 hours. Long term indicates treatment with drug at 1 nM for 2 weeks. Log-fold change is calculated from unstimulated/untreated condition. Decrease in MEK phosphorylation post trametinib treatment is observed between 1-6 hours.

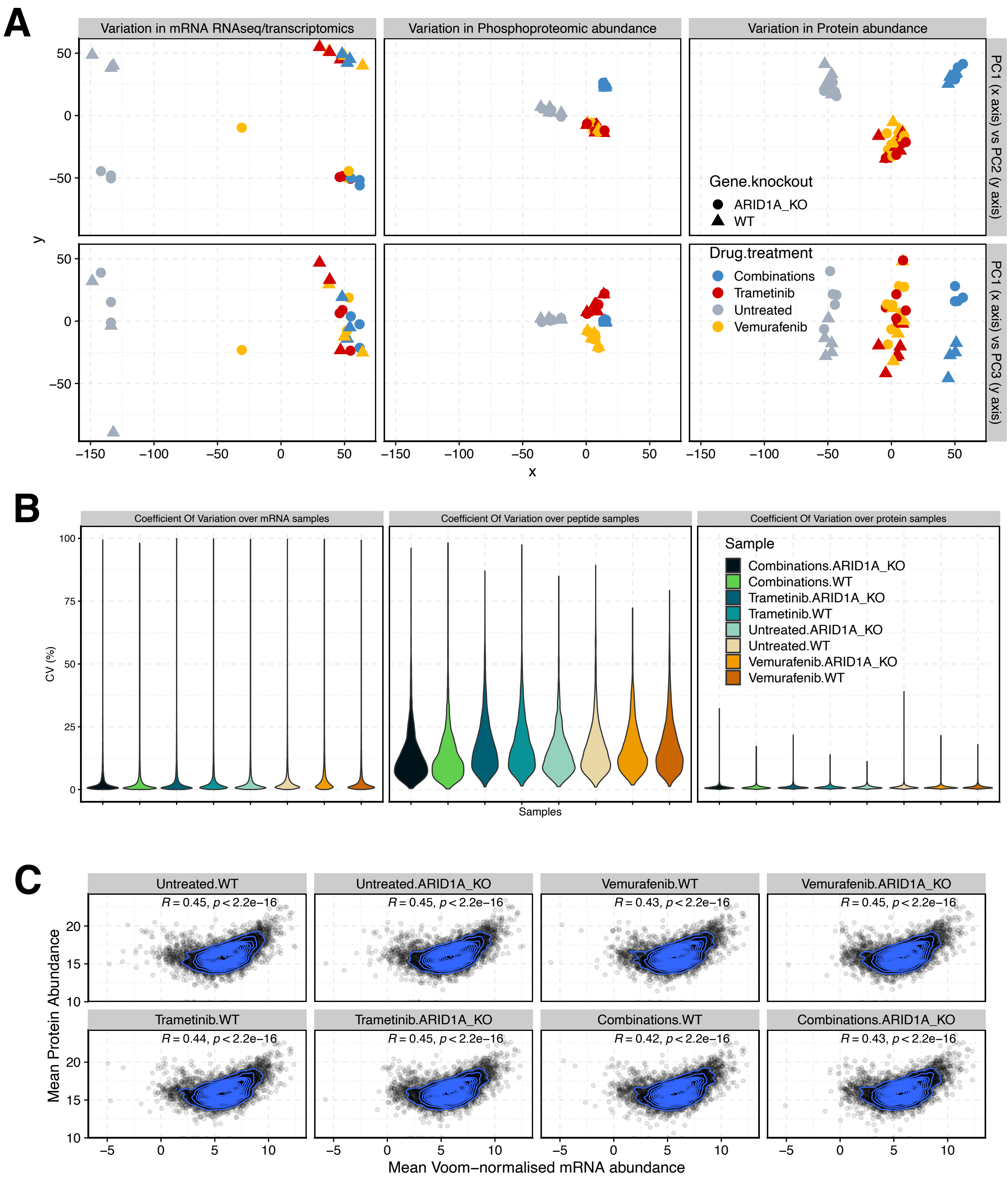


**Supplementary Figure S2. Summary of count information for ’omics data in different experimental conditions. A.** Multidimensional scaling (MDS) for RNAseq (left), peptides (middle) and proteins (right) showing the genetic background (shape) and the drug treatment (colour) experimental conditions. **B.** Coefficient Of Variation (COV) plots for RNAseq (left), peptides (middle) and proteins (right).**C.** Comparison of protein abundance (y axis), versus mRNA abundance (x axis) for different experimental conditions.

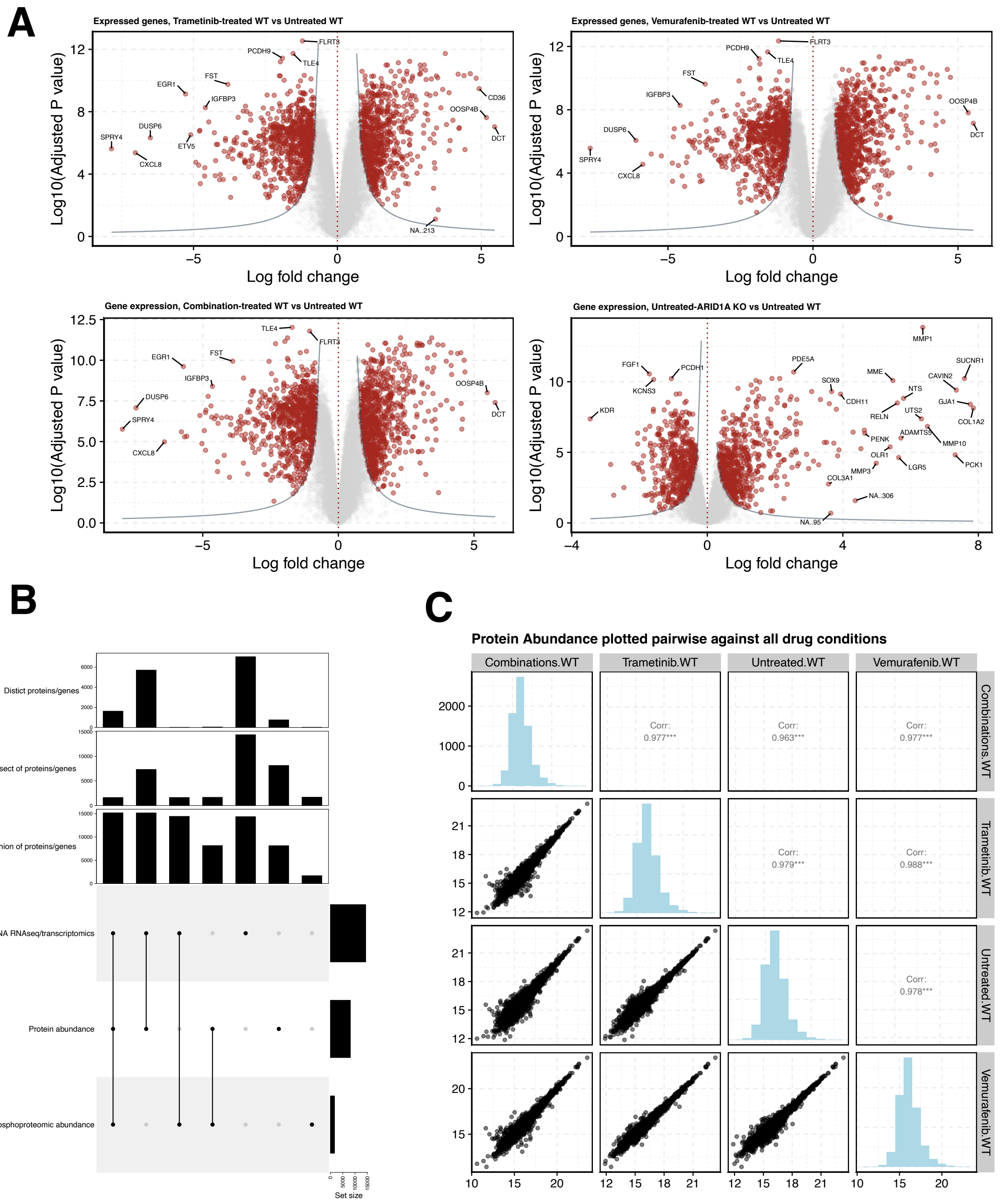


**Supplementary Figure S3. Differential expression analysis and quantified proteins/genes across different ’omics data. A.** Volcano plots for abundant gene transcripts showing significance (FDR-adjusted p value, y axis) and log-fold change (x axis). LFCs for this data as well as phosphopeptides and proteins are available in the supplementary tables. **B.** Upset plot showing the extent of distinct proteins, genes and peptides quantified from each ’omics experiment. Barplots show the number of readings counted in each intersection of the data. **C.** Plots showing the relation between protein abundance for isolated drug treatments.

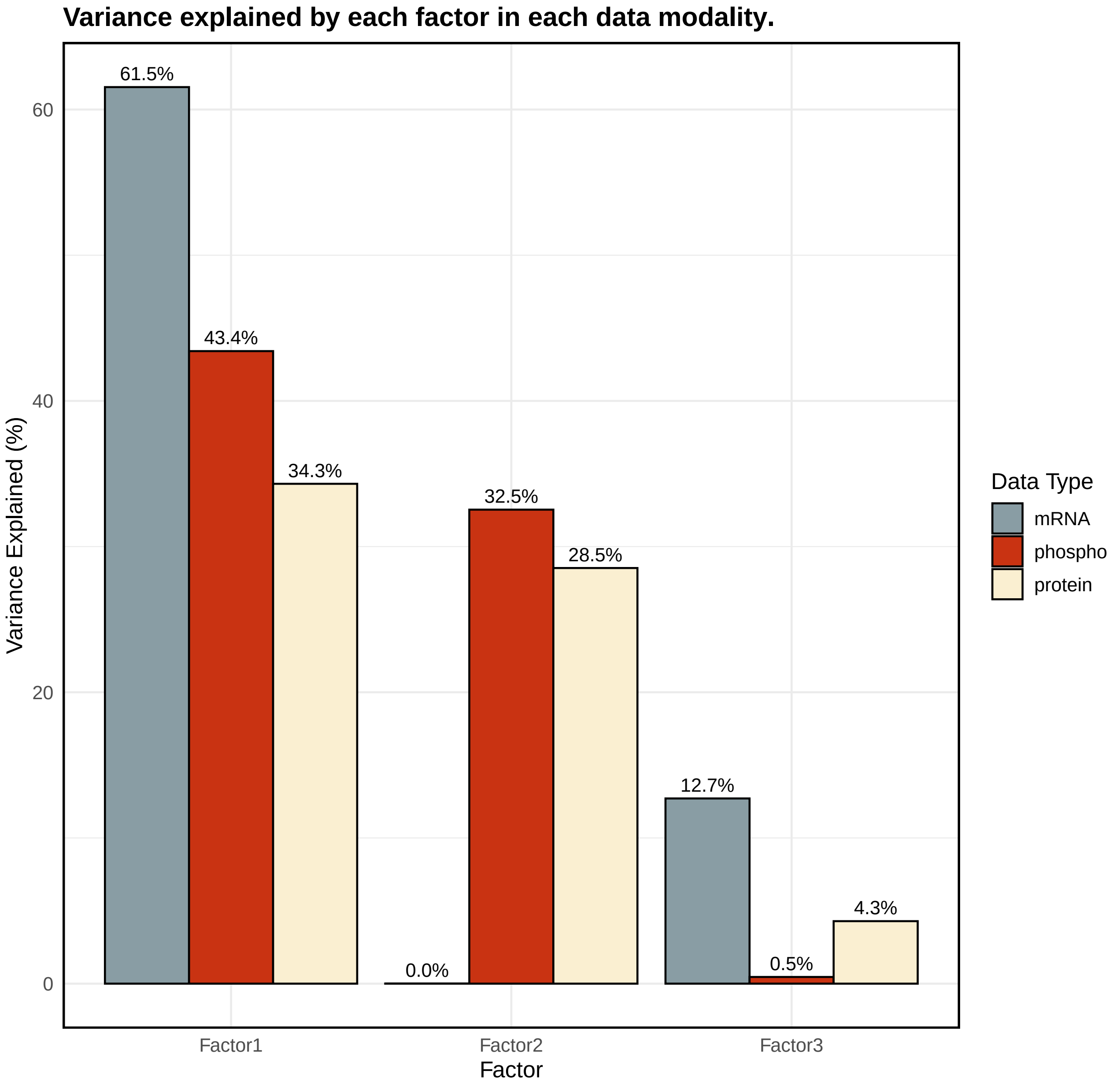


**Supplementary Figure S4. Variance decomposition derived from MOFA analysis.** Barplot showing the variance decomposition, assessing the proportion of variance explained by each factor in each 'omics modality

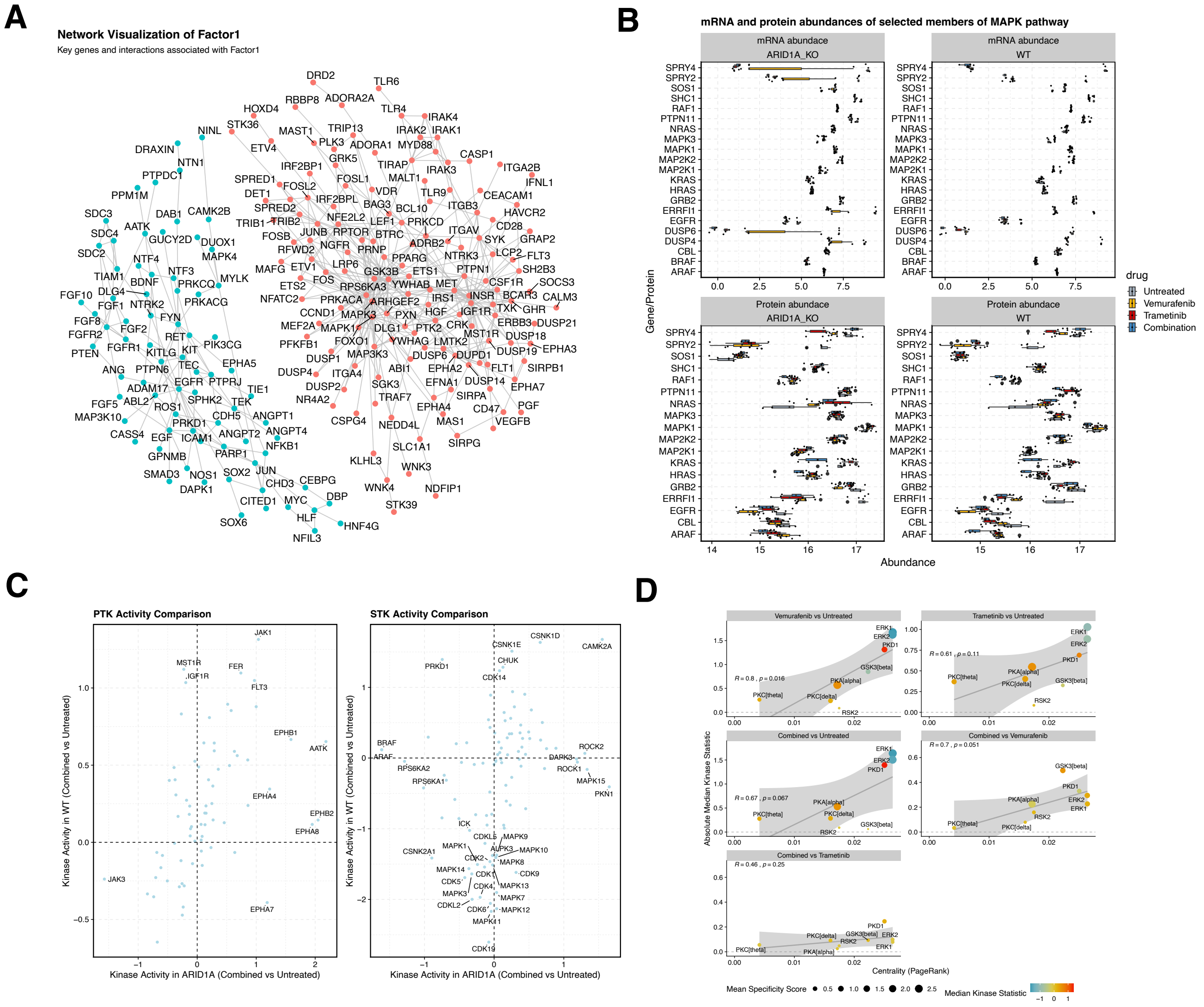


**Supplementary Figure S5. Drug-agnostic network-based changes. A.** PhueGO-derived Network underlying changes associated with Factor1. Nodes represent proteins and the edges between them represent known interactions between those proteins. Red indicates networks associated with upregulated processes and blue indicates networks associated with downregulated processes. **B.** Changes in abundance (x axis) of MAPK negative feedback regulators (y axis) identified by Gerosa *et al.*. Colour represents the drugs used. **C.** Kinase activity assay showing the activation of kinases after combined drug treatment in ARID1A KO (x axis), vs parental (y axis) A375 cell lines.**D.** Absolute values for median kinase activity correlated with centrality of network in S5A.

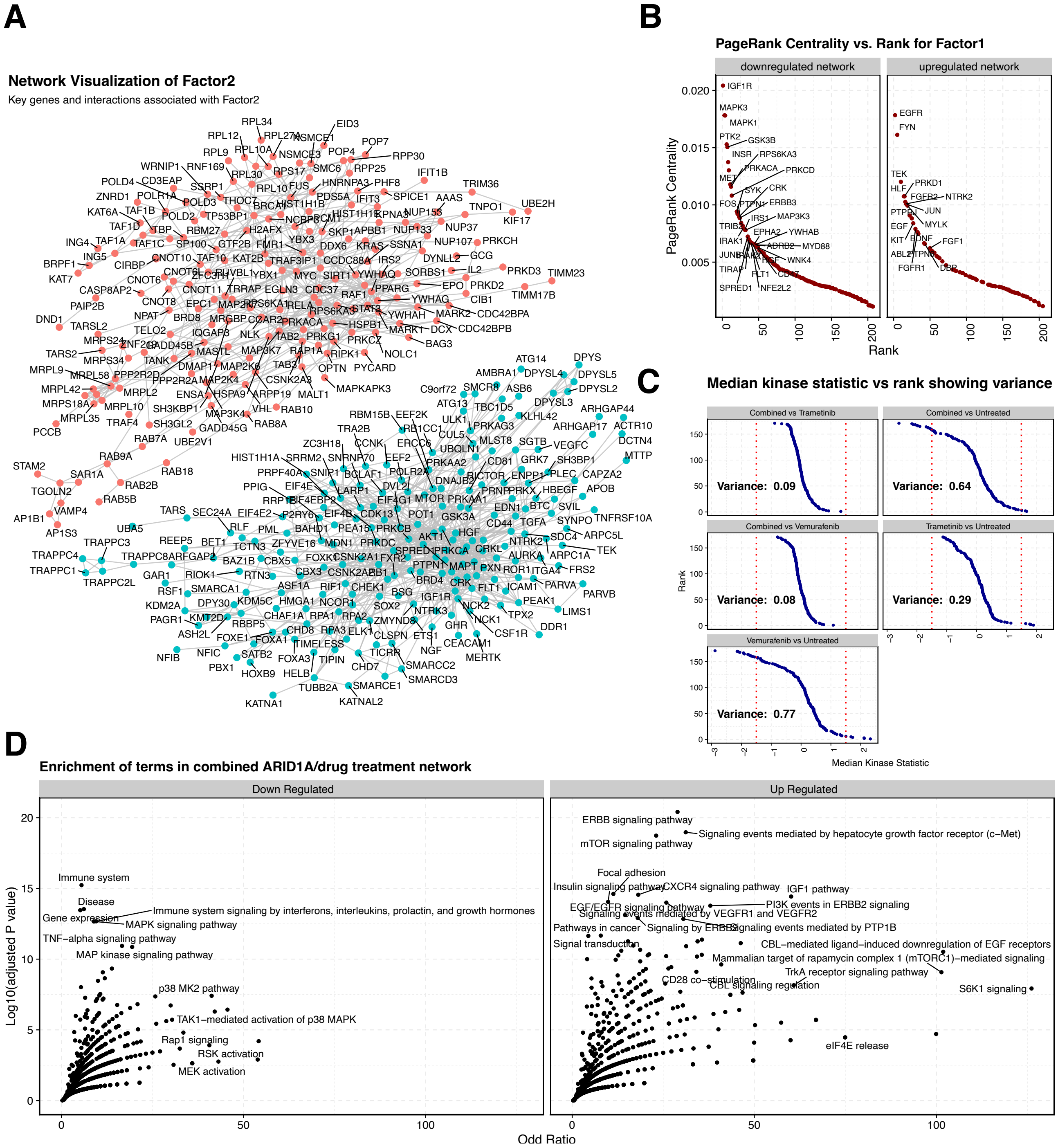


**Supplementary Figure S5. Combination therapy-specific network-based changes. A.** PhueGO-derived Network underlying changes associated with Factor2. Nodes represent proteins and the edges between them represent known interactions between those proteins. Red indicates networks associated with upregulated processes and blue indicates networks associated with downregulated processes. **B.** Centrality (PageRank, y axis) of nodes in the network represented in Figure S6A, ranked by order of size (x axis). **C.** Changes in median kinase activity (x axis), showing the difference in variance amongst all experimental conditions. **D.** Functional enrichment analysis of upregulated and downregulated genes in Factor2 derived network, using the pathway database Bioplanet 2019. Axis represent strength of the association (Odds ratio, x axis) and the significance (Log10(FDR adjusted value), y axis).

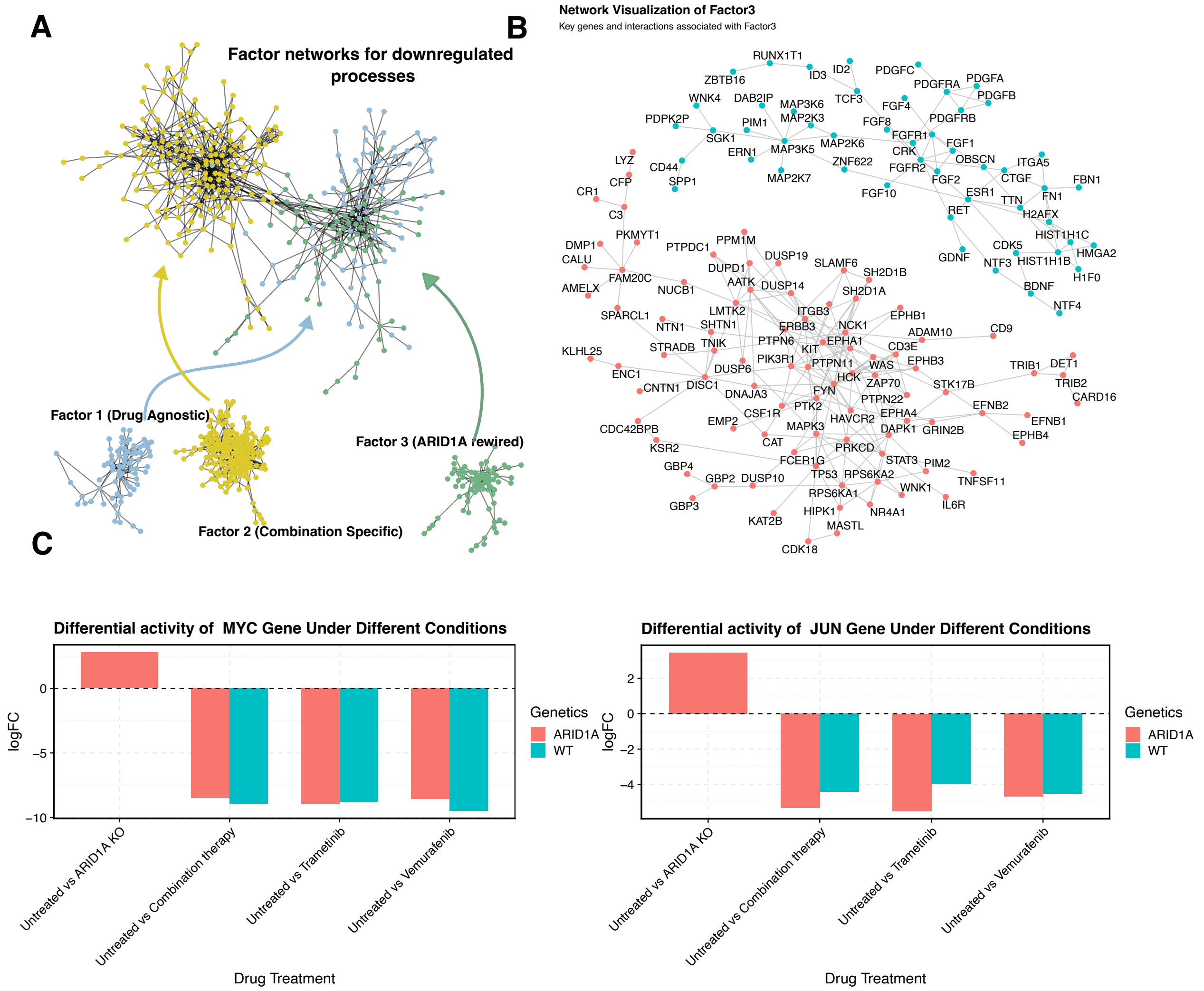


**Supplementary Figure S7. ARID1A-rewired network-based changes. A.**Graph showing the increased connectivity of nodes from both Factor 1 (blue) and Factor 3 (green) compared to Factor 2 (yellow). **B.** PhueGO-derived Network underlying changes associated with Factor3. Nodes represent proteins and the edges between them represent known interactions between those proteins. Red indicates networks associated with upregulated processes and blue indicates networks associated with downregulated processes. **C.** Plot showing the change in activation of TFs MYC (left) and JUN (right), based on the relative abundance of their known regulons as calculated using the ColectTRI database. This shows the difference in condition between ARID1A KO A375 cell lines (red) and parental cell lines (blue).

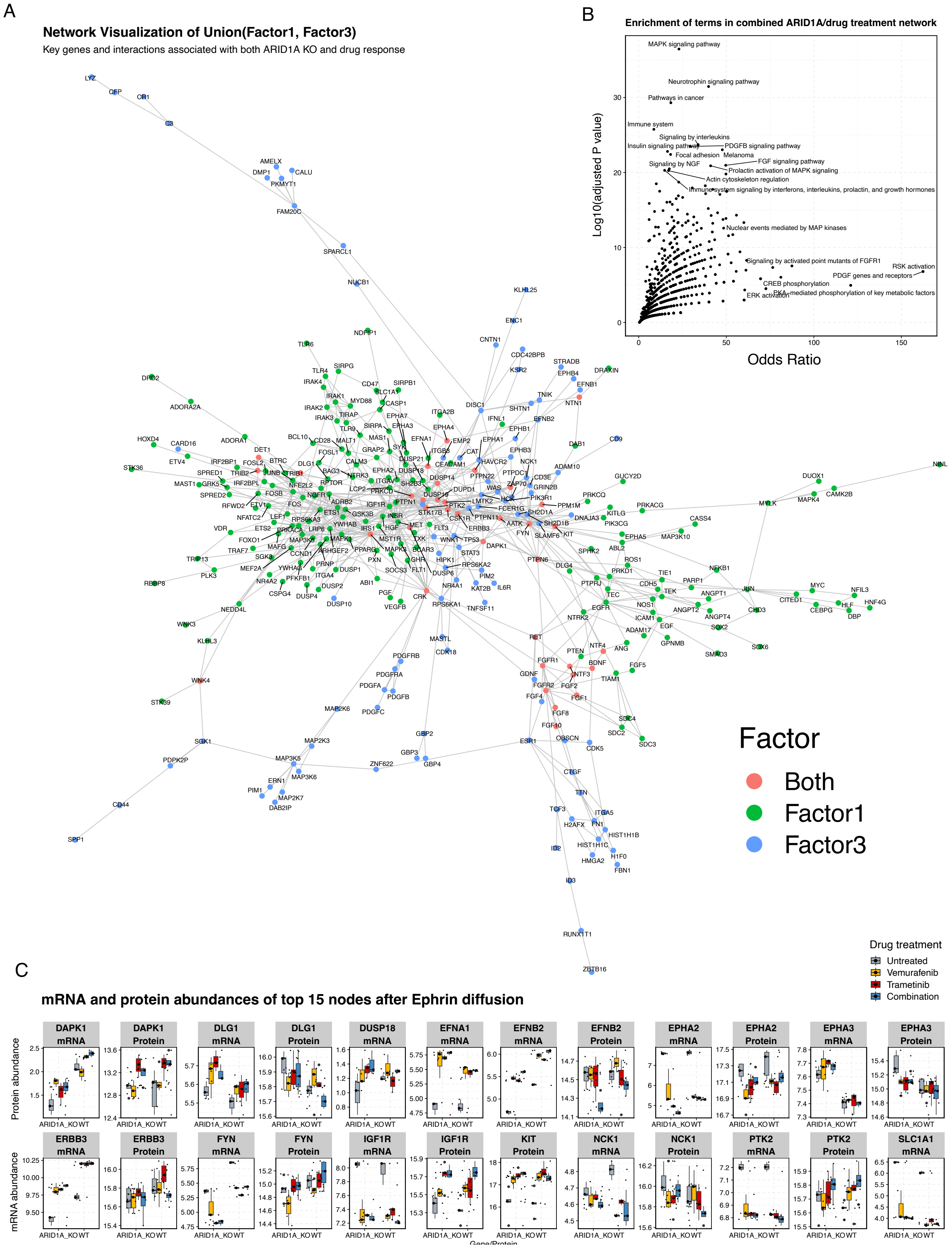


**Supplementary Figure S8. Cross-talk in network-based changes between ARID1A-rewired and drug-agnostic response. A.** Combined network from PhueGO-derived Network underlying changes associated with Factor3 (blue) and Factor1 (green). Nodes represent proteins and the edges between them represent known interactions between those proteins. **B.** Functional enrichment analysis of genes in combined Factor1/3 network, using the pathway database Bioplanet 2019. Axis represent strength of the association (Odds ratio, x axis) and the significance (Log10(FDR adjusted value), y axis). **C.** Changes in abundance (x axis) of downstream effectors of Ephrin (y axis) identified by maximum flow calculations through the combined Factor1/3 network. Colour represents the drugs used.

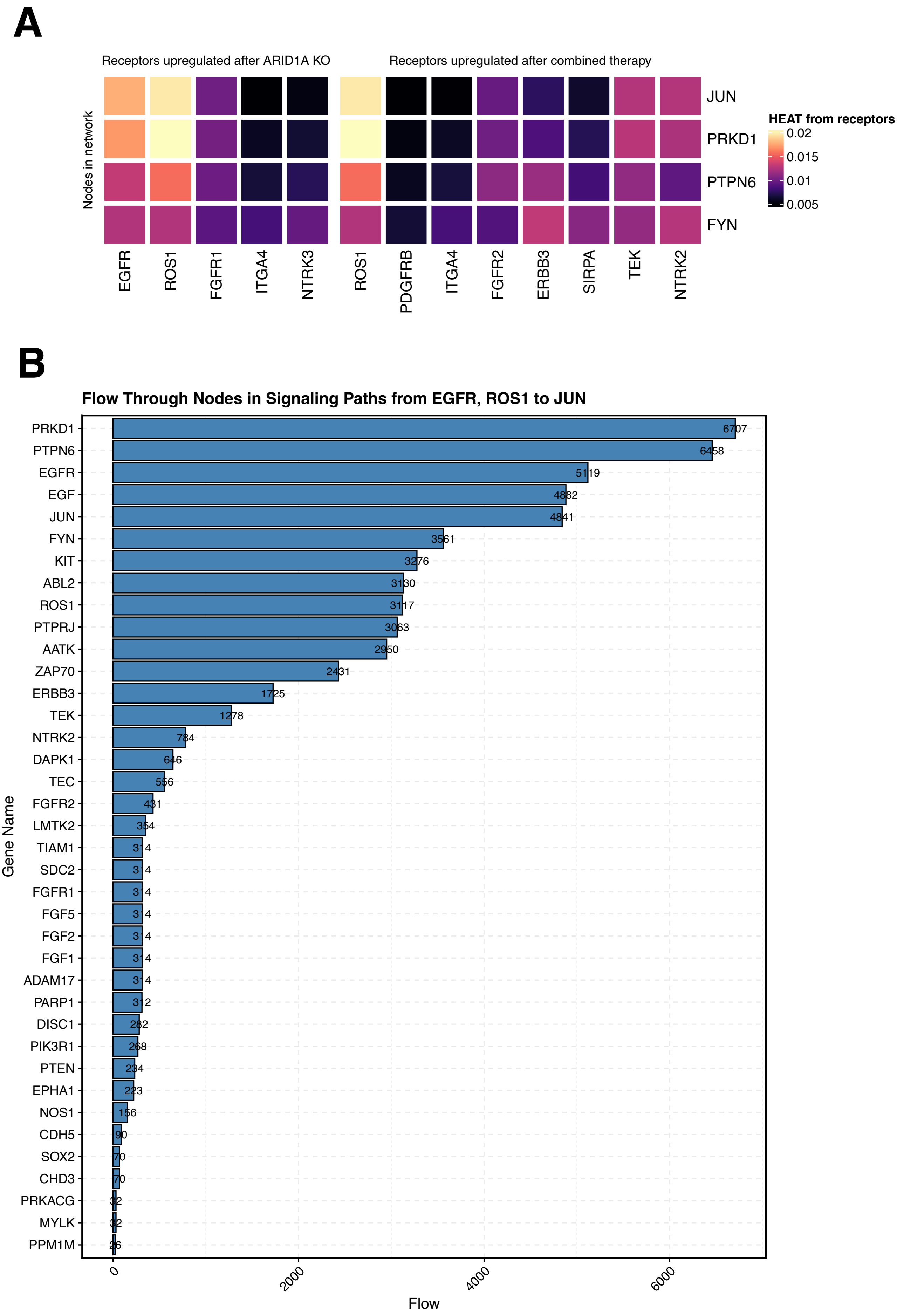


**Supplementary Figure S9. Network propagation illustrating the affect of dysregulated receptors. A.** Heatmap showing network propagation from individual receptors (columns) and their propagation to selected nodes (rows). **B.** Barplot showing the flow (x axis) through nodes (y axis) within the combined Factor1/3 network from EGFR and ROS1 to JUN, ranked in order of flow.

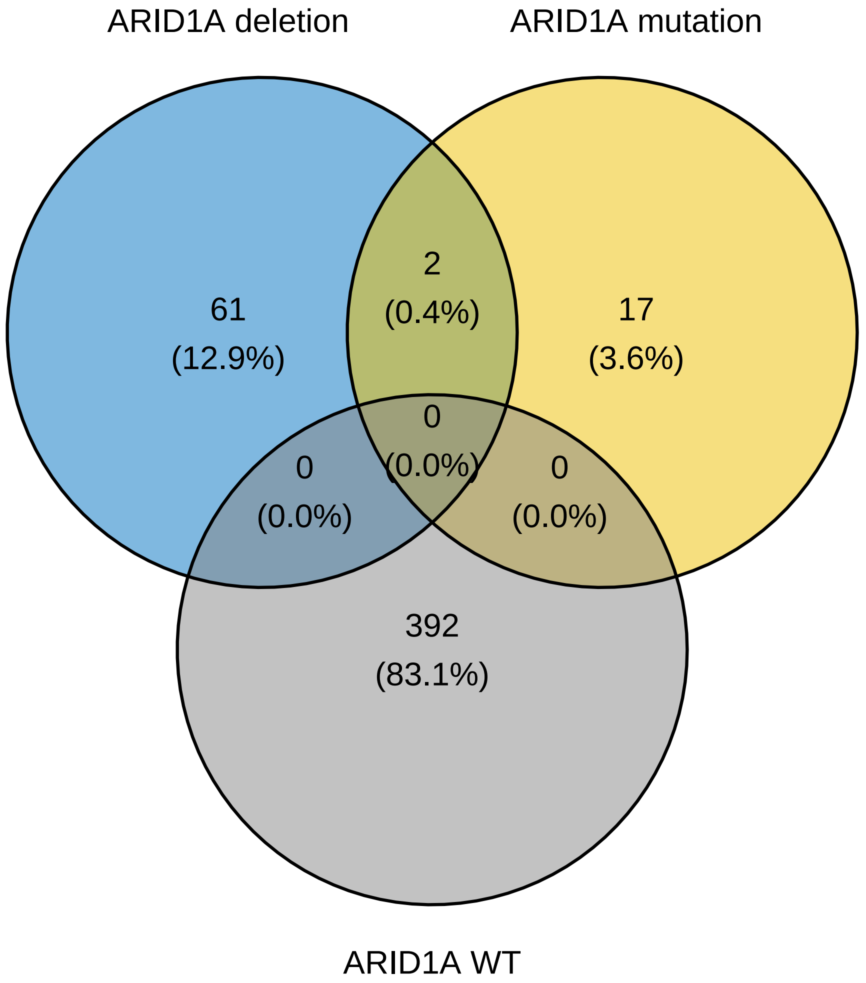


**Supplementary Figure S10. Summary of melanoma patients extracted from TCGA.** Venn diagrams showing patients stratified by whether they have an ARID1A mutation, deletion or neither.
